# Post-Transcriptional Circadian Regulation in Macrophages Organizes Temporally Distinct Immunometabolic States

**DOI:** 10.1101/2020.02.28.970715

**Authors:** Emily J Collins, Mariana P Cervantes-Silva, George A Timmons, James R O’Siorain, Annie M Curtis, Jennifer M Hurley

## Abstract

Our core timekeeping mechanism, the circadian clock, regulates an astonishing amount of cellular physiology and behavior, playing a vital role in organismal fitness. While the mechanics of circadian control over cellular regulation can in part be explained by the transcriptional activation stemming from the positive arm of the clock’s transcription-translation negative feedback loop, research has shown that extensive circadian regulation occurs beyond transcriptional activation in fungal species and data suggest that this post-transcriptional regulation may also be preserved in mammals. To determine the extent to which circadian output is regulated post-transcriptionally in mammalian cells, we comprehensively profiled the transcriptome and proteome of murine bone marrow-derived macrophages in a high resolution, sample rich time course. We found that only 15% of the circadian proteome had corresponding oscillating mRNA and this regulation was cell intrinsic. Ontological analysis of oscillating proteins revealed robust temporal enrichment for protein degradation and translation, providing potential insights into the source of this extensive post-transcriptional regulation. We noted post-transcriptional temporal-gating across a number of connected metabolic pathways. This temporal metabolic regulation further corresponded with rhythms we observed in ATP production, mitochondrial morphology, and phagocytosis. With the strong interconnection between cellular metabolic states and macrophage phenotypes/responses, our work demonstrates that post-transcriptional circadian regulation in macrophages is broadly utilized as a tool to confer time-dependent immune function and responses. As macrophages coordinate many immunological and inflammatory functions, an understanding of this regulation provides a framework to determine the impact of circadian regulation on a wide array of disease pathologies.

## INTRODUCTION

Much of life on earth is exposed to the regular day/night cycle of the planet and has evolved circadian rhythms to anticipate these predictable changes (Eckel-Mahan and Sassone-Corsi, 2013). Circadian rhythms are generated by a molecular oscillator, or clock, which adjusts innumerable cellular functions to regulate behaviors, encompassing everything from luminescence in fungi to sleep in humans (Dunlap, 1999; Patke, Young and Axelrod, 2019). The canonical view of physiological circadian regulation is via a transcription-translation based negative-feedback loop, where the heterodimeric transcription factor positive-arm complex regulates the creation of both the negative arm as well as a host of other genes not involved in the feedback loop, termed clock-controlled genes (*ccgs*). These oscillations in transcript levels are assumed to translate to oscillations in protein levels (Hurley, Loros and Dunlap, 2016; Takahashi, 2016). Utilizing many different organisms and tissue types, transcriptional studies of clock output have found that large proportions of the transcriptome oscillate in a circadian manner and that the transcripts regulated by the circadian clock are highly tissue-specific (Hughes *et al.*, 2009; Abruzzi *et al.*, 2011; Zhang *et al.*, 2014; Mure *et al.*, 2018).

However, recent evidence has established that post-transcriptional and post-translational mechanisms play major roles in regulatory output from the circadian clock, resulting in circadianly oscillating proteins that are not derived from oscillating transcripts and vice versa (Reddy *et al.*, 2006; Kojima, Shingle and Green, 2011; Chiang *et al.*, 2014; Hurley *et al.*, 2014, 2018; Mauvoisin *et al.*, 2014; Robles, Cox and Mann, 2014; J. Wang *et al.*, 2017; Green, 2018; Wang *et al.*, 2018; Mauvoisin and Gachon, 2019). Despite this mounting evidence, there is a lack of studies that utilize adequate sampling depth, resolution, and time course length to generate highly-powered datasets that can probe the global regulatory relationships between the circadian transcriptome and proteome. To determine if the large extent of post-transcriptional regulation was conserved in mammalian cells, we analyzed the global transcriptome and proteome of murine Bone Marrow-Derived Macrophages (BMDMs) every two hours, over 48 hours, in triplicate using RNA-seq and Tandem Mass Tag Mass Spectrometry (TMT-MS) respectively.

BMDMs were selected for this in-depth investigation as previous work has shown that macrophages possess robust circadian clocks and demonstrate circadian control of their key immune functions, such as phagocytosis, cytokine release, differentiation and migration in/out of various tissue compartments (Bourin *et al.*, 2002; Hayashi, Shimba and Tezuka, 2007; Keller *et al.*, 2009; Gibbs *et al.*, 2011; Sato *et al.*, 2014; Wang *et al.*, 2016; Kiessling *et al.*, 2017; Silver *et al.*, 2018; Pick *et al.*, 2019). Macrophages play a vital role in the initiation, sustainment, and resolution of both acute and chronic inflammation, and it has been suggested that the disruption of these circadian rhythms in immunity and inflammation may play a critical part in the negative impact of clock disruption on human health (Bechtold, Gibbs and Loudon, 2010; Arjona *et al.*, 2012; Cermakian *et al.*, 2013; Evans and Davidson, 2013; Scheiermann *et al.*, 2018)(Curtis *et al.*, 2014; Geiger, Fagundes and Siegel, 2015; Labrecque and Cermakian, 2015; Scheiermann *et al.*, 2018). Finally, as macrophages live for months at a time, circulating in the bloodstream before long in-tissue residencies, the influence of their circadian clocks would be of more consequence than shorter-lived immune cells (van Furth and Cohn, 1968). However, much is left to be desired in the study of macrophage rhythms as most prior studies have used transcriptional and ex/in vivo methods to determine macrophage output pathways, complicating what rhythms are generated by the presence of rhythmic systemic cues vs what is intrinsically controlled by macrophage clocks.

When we compared the oscillating transcriptome to the oscillating proteome, we found that the extent of post-transcriptional regulation in macrophages is greater than in any previously studied system, with only 15% of oscillating proteins pairing with an oscillating mRNA. Gene ontological analysis of these oscillating proteins suggested that this post-transcriptional regulation, unlike previously studied cell types, stems both from the regulation of translation as well as degradation (Hurley *et al.*, 2018). In addition, we found that metabolism, particularly the process of ATP generation, was highly post-transcriptionally regulated. Evidence of post-transcriptional metabolic regulation extended beyond changes in abundance of central metabolic complexes, as we also noted oscillations in key proteins involved in mitochondrial morphology, a central regulator of macrophage energy production and the immune response (Langston, Shibata and Horng, 2017; Y. Wang *et al.*, 2017). When we analyzed the behaviors controlled by these post-transcriptionally regulated proteins in live cells, we confirmed corresponding oscillations in ATP generation and mitochondrial morphology. We also investigated phagocytosis in our macrophages and found that this process aligned with the time of day predicted by the metabolic state, suggesting that post-transcriptional circadian regulation is essential in timing metabolic control of macrophages to regulate the immune response.

## RESULTS

### Analysis of the Macrophage Circadian Transcriptome and Proteome Demonstrates Extensive Circadian Regulation

To determine if the substantial, physiologically relevant, circadian post-transcriptional regulation noted in the fungal system was conserved in higher eukaryotes, macrophages were chosen as a model as they have previously-described robust circadian oscillations and play an essential role in the immune system (Hayashi, Shimba and Tezuka, 2007; Keller *et al.*, 2009; Silver *et al.*, 2012; Sato *et al.*, 2014; Labrecque and Cermakian, 2015). Bone-Marrow Derived Macrophages (BMDMs) were derived from bone marrow progenitor cells extracted from Per2::Luc C57/BLK6 mice and differentiated with recombinant M-CSF (Yoo *et al.*, 2004). Flow cytometry confirmed that >99% of cells displayed the macrophage-specific markers CD11b and F4/80, demonstrating complete differentiation into macrophages (Supplemental Figure 1A) (Zhang, Goncalves and Mosser, 2008). We extracted total RNA and protein from these serum-shock-synchronized BMDMs every 2 hours over 48 hours in triplicate, totaling 6 independent time courses (Figure 1A, Supplemental Figure 1B). Luminescence traces were used to confirmed that our protocol resulted in reliable, ∼24-hour PER2 oscillations (Supplemental Figure 1C)

**Figure 1.**
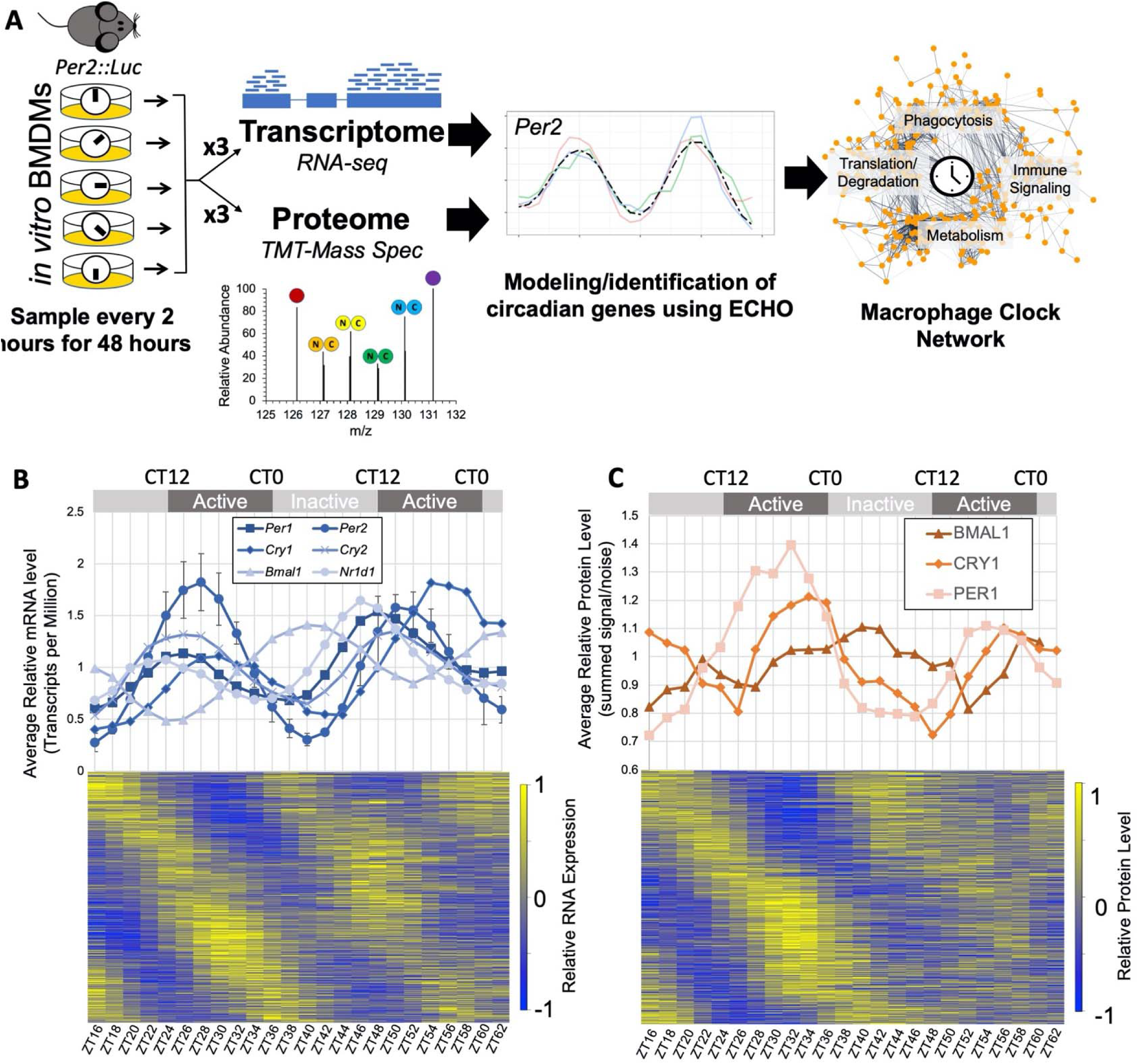
Multi-Omics Profiling Details Extensive Circadian Regulation of the Macrophage Transcriptome and Proteome. (A) A schematic of the analysis of macrophage circadian regulation. Bone marrow derived macrophages (BMDMs) from Per2::Luc mice were synchronized *in vitro* via serum shock and sampled in triplicate for global proteome and transcriptome profiling with 10-plex Tandem Mass Tag mass spectrometry and RNA-seq respectively. Circadianly-oscillating genes were identified using the ECHO program to determine the macrophage circadian protein-protein interaction network. (B) All detected clock gene transcripts oscillated circadianly and error bars show the standard deviation for *Per2* as a representative gene for the variation between triplicate values. The circadian time (CT) and inactive/active phases were inferred from comparison of *Per2* oscillation timing we observed to the oscillation of *Per2* in (Keller *et al.*, 2009). Heatmap in bottom panel shows relative expression for all identified circadian transcripts. (C) All detected clock proteins exhibited circadian abundances with delay from their respective transcripts. Points shown are an average of the summed signal/noise of peptides for the identified protein for up to three replicates depending on the detection at the given time point. Heatmap in bottom panel shows relative expression for all identified circadian proteins.

Quantitative RNA-sequencing was performed to analyze the transcriptome. In précis, samples were ribo-depleted before library creation and Illumina 10X total RNA-sequencing, so as to quantify both coding and non-coding RNA expression (see methods). An ideal read depth of an average of 74.9 million reads per sample was achieved (Supplemental File 1-will be published upon peer review). Read alignments for each gene were normalized for varying gene size by converting to Transcripts per Million. Quantitative proteomic analysis was completed using Tandem Mass Tag – Mass Spectrometry. Protein samples were isobarically-mass tagged after digestion and analyzed utilizing eight multiplex groups. Each multiplex contained nine randomized samples to prevent biased batch effects and one sample of proteins pooled from all samples was run in all eight multiplex sets to assist in relative quantitation between multiplexes (see methods). Across all multiplexes, 162,920 unique peptides were identified in at least one time point, comprising a total of 10,162 distinct proteins detected, with 6,290 proteins detected in every time point (Supplemental File 2-will be published upon peer review).

The transcriptome and proteome data were pre-processed and analyzed using the previously published tools LIMBR and ECHO, designed for circadian time course analysis (De los Santos *et al.*, 2017, 2020; Crowell *et al.*, 2019). Any transcript/peptide with <70% detection across all 75 time points was disqualified from our investigation, after which any missing values were imputed and underlying batch effects removed using the LIMBR pipeline (Crowell *et al.*, 2018). After LIMBR pre-processing, a total of 6,050 proteins and 36,451 RNAs qualified for further analysis. These transcripts and proteins were categorized as oscillating/not-oscillating using the ECHO program in the free-run mode (De los Santos *et al.*, 2020).

Once we identified our oscillating proteins and transcripts, we examined both datasets for clock gene levels. We found that each identified positive- and negative-arm clock gene known to oscillate in a circadian manner did so at both the RNA (Figure 1B) and protein levels (Figure 1C). Clock gene transcripts including *Arntl, Clock, Cry1, Cry2, Nr1d1, Per1* and *Per2* were identified as circadian with low variation between replicates (Figure 1B, top panel & Supplemental Figure 2A). Due to known low abundance levels and our stringent detection cutoffs, no clock genes were classified as detected in the proteome. However, manual examination of pre-LIMBR curated average levels of clock gene proteins at each time point demonstrated oscillations of both positive (BMAL1) and negative arm (PER1, CRY1) clock proteins (Figure 1C, top panel). As predicted, the positive and negative arm protein oscillations were anti-phasic and a few hours delayed relative to their associated transcripts (Figures 1B & 1C).

Based on the ECHO-modeled periods for known clock gene transcripts, genes were considered to be circadianly oscillating if Benjamini-Hochberg adjusted p-value was <0.05, the amplitude change coefficient was between +/- 0.15 and the period between 20-28 hours. This classified 5,790 transcripts as circadian, 15.8% of the detected macrophage transcriptome (Figure 1B). A similar proportion, 15.3%, of coding transcripts were circadian (n= 3,160) within the coding-only detected transcriptome (n=20,699). When analyzing the distributions of modeled parameters, we noted a bias towards a shorter period length (between 20-22 hrs) and that most oscillations decreased in amplitude (damped) over time (Supplemental Figure 2B) (De los Santos *et al.*, 2017, 2020). For ECHO-modeled protein periods, we broadened the period range of what we considered circadian to 18-30 hrs. to accommodate higher levels of technical noise attributable to proteomics-based techniques (Hurley *et al.*, 2018). Analysis of the proteome identified 1,778 proteins as circadian, meaning 29% of the detected proteome is under circadian control (Figure 1C). There was little bias in period length detected and more widespread damping over time as compared to the transcriptome (Supplemental Figure 2B). In both cases, the vast majority of genes in the circadian RNA and protein sets had p-values much lower than 0.01, indicating statistically accurate modeling of these genes by ECHO (Supplemental Figure 2B, lower panel).

### Post-Transcriptional and Post-Translational Control in the Late Active Phase Contributes to Circadian Output

In an exhaustive study, Mure et al. concluded that there was extensive circadian gating on mammalian transcripts, but did not investigate the impact of these transcriptional waves on protein levels and, by extension, cell physiology (Mure *et al.*, 2018). We therefore probed our datasets for circadianly-gated timing of the peak phases of RNA and protein and found that macrophage circadian RNA and protein levels both peak in two “waves,” one smaller wave in the late inactive phase (CT3-15, Circadian RNAs n=2,724, Circadian Proteins n= 807), and one larger wave in the late active phase (CT15-3, Circadian RNAs n= 3,066, Circadian Proteins n= 947) (Figure 2A). When we subsetted our RNA data to only include circadian RNAs that encoded proteins (n=2,801), the RNA waves became more evident, occurring ∼2 hours before the corresponding translational wave (CT1-13 Circadian RNAs n=1,415, CT13-1 Circadian RNAs n=1,386) (Figure 2A).

**Figure 2.**
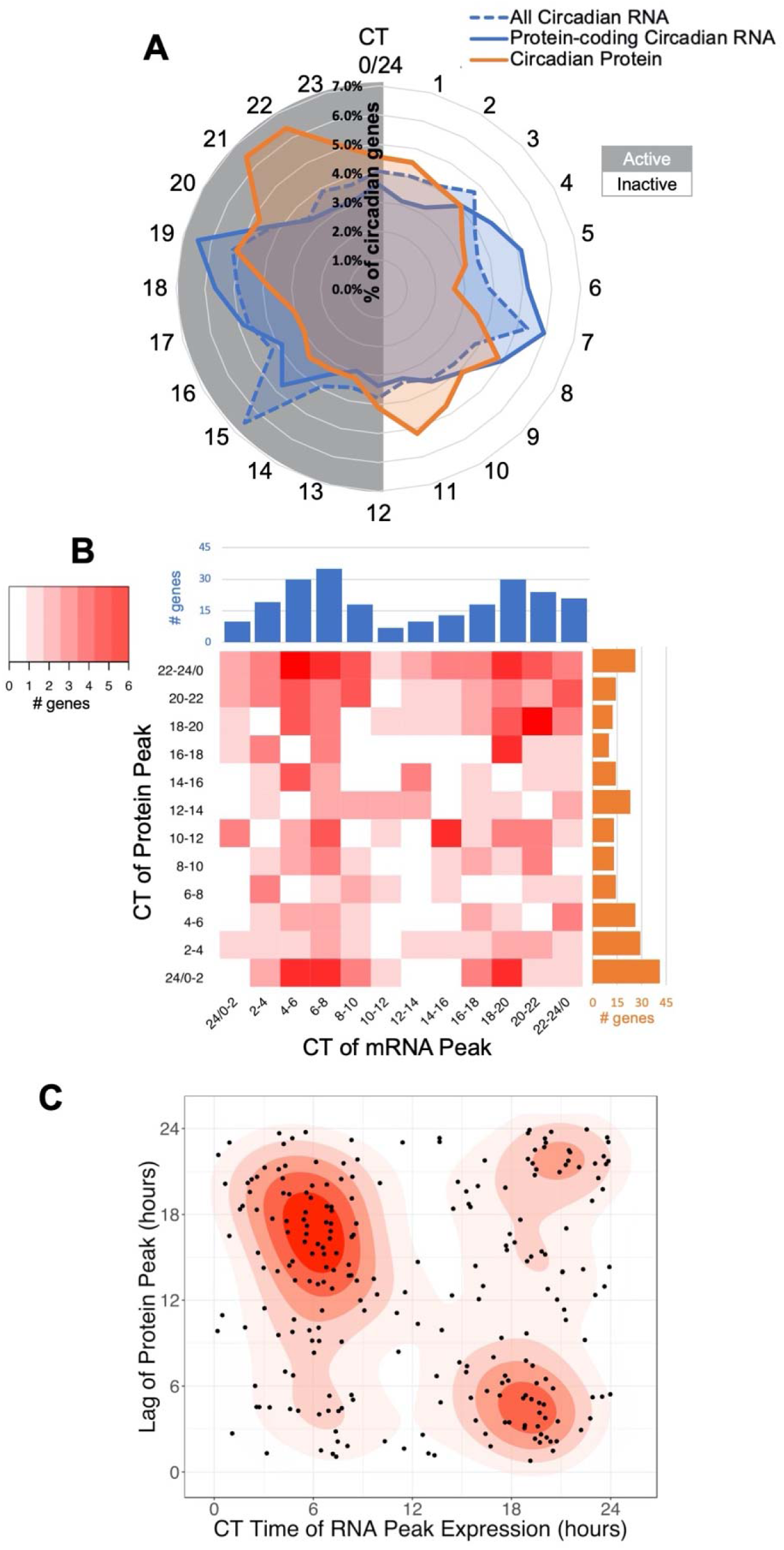
Comparison of Circadian Transcripts and Proteins Reveals A Significant Role for Post-Transcriptional and Post-Translational Regulation. (A) The percentage of total oscillating transcripts (blue) or proteins (orange) peaking at a given time was plotted on a radial histogram by Circadian Time (CT) of peak, binned in 1 hour windows. (B) A heat map comparing the difference in the peak phases of corresponding oscillating transcripts (blue) and proteins (orange) binned into 2 hour intervals over circadian time for genes circadian at both the mRNA and protein level. (C) Density graph displaying the lag time to protein creation from the peak time of corresponding oscillating transcript levels for genes circadian at both the mRNA and protein level.

The fixed delay between RNA and protein peaks suggested a correlation between the temporally-related RNA and protein waves, i.e. mRNAs are transcribed and shortly after translated into proteins, as would be suggested by the canonical circadian positive-arm regulation framework. To determine if these general temporal waves aligned with specific protein/RNA relationships, we subsetted our datasets to examine only genes that were detected and rhythmic in both the transcriptome and proteome. This identified 250 genes that were circadian at both the mRNA and protein level (Supplemental Figure 3A) (Supplemental File 3-will be published upon peer review). We compared the peak time of each mRNA with the peak time of its corresponding protein and found a surprisingly low correlation between the RNA and protein pairs. While the majority of oscillating proteins with oscillating mRNA peaked during the late active wave of translation, the peak phase of their corresponding mRNAs was evenly distributed between the two transcriptional waves (Figure 2B). The dominance of this late active wave peak for protein abundance appeared to be specific to proteins with associated oscillating mRNA, as the peak level timings of proteins that did not have oscillating mRNA were more evenly distributed (Supplemental Figures 3B & 3C). We also noted that the time of day that a protein reached its peak levels correlated with the phase delay between peak RNA and protein levels (Figure 2C). Transcripts that peaked near CT6 took 15-18 hours to reach their respective protein peak, whereas transcripts peaking around CT18 only took 3-6 hours to reach their protein peak. This suggested that translation of transcripts occurs more readily during the late active phase, implying temporal differences in post-transcriptional regulation.

We then expanded our analysis to investigate the overall relationship between oscillating RNAs and proteins. We found that while our transcriptome largely encompassed our proteome (n= 5,833 total genes), there was little overlap between circadian transcripts and circadian proteins (Supplemental Figures 3A & 3D). 68% of circadian transcripts did not result in a circadian protein and 86% of circadian proteins did not derive from a circadian transcript. As this discrepancy was considerably higher than has been previously reported, we wanted to rule out that this lack of overlap was due to the more lenient period cutoffs used for proteins to be considered circadian (Hurley *et al.*, 2018). We therefore evaluated the overlap using the more or less lenient period cutoffs for both datasets (Supplemental Figures 3E & F) and found that the ratios remained skewed towards minority overlap, indicating strong evidence for post-transcriptional regulation in these BMDMs.

### Post-Transcriptional and Post-Translational Mechanisms are Abundant in the Late Active Phase

The extensive evidence of post-transcriptional circadian control described above led us to examine clock-timed biological processes that could contribute to this regulation. Past studies have shown that translation may be the key process underlying circadian post-transcriptional regulation and gene ontological enrichment analysis to examine the 250 genes rhythmic at both the mRNA and protein levels found that they were enriched for processes involved in translation (Bonferroni P-value = 2.72E-4, Fold enrichment = 5.18) (Supplemental Table 1) (Ashburner *et al.*, 2000; Hurley *et al.*, 2014, 2018; Caster *et al.*, 2016; Fabregat *et al.*, 2018; Mi *et al.*, 2019). Therefore, we examined the circadian proteome to determine if a significant proportion of the clock output has functionality in translation. First, we split the proteins into two groups in line with the temporal “waves” observed in Figure 2A. Proteins that peaked in abundance between CT3 and CT15 were considered “late inactive wave” proteins and proteins that peak between CT15 and CT3 were considered “late active wave” proteins. Both categories were assessed for enrichments for Reactome categories using Amigo/Panther enrichment analysis tools as well as interrogated to determine protein-protein interactions (Ashburner *et al.*, 2000; Fabregat *et al.*, 2018; Mi *et al.*, 2019). Proteins belonging to significantly enriched (Bonferroni adjusted p-value <0.05) Reactome categories were color coded according to their category and graphed by their protein-protein interactions with String-DB (Figure 3, Supplemental Figure 4 & 5) (Szklarczyk *et al.*, 2014). We further utilized Phase Set Enrichment Analysis (PSEA) with curated murine gene ontology gene sets to investigate protein enrichment by time of day (Supplemental Figure 6A) (Zhang *et al.*, 2016; Bares and Ge, 2019).

**Figure 3.**
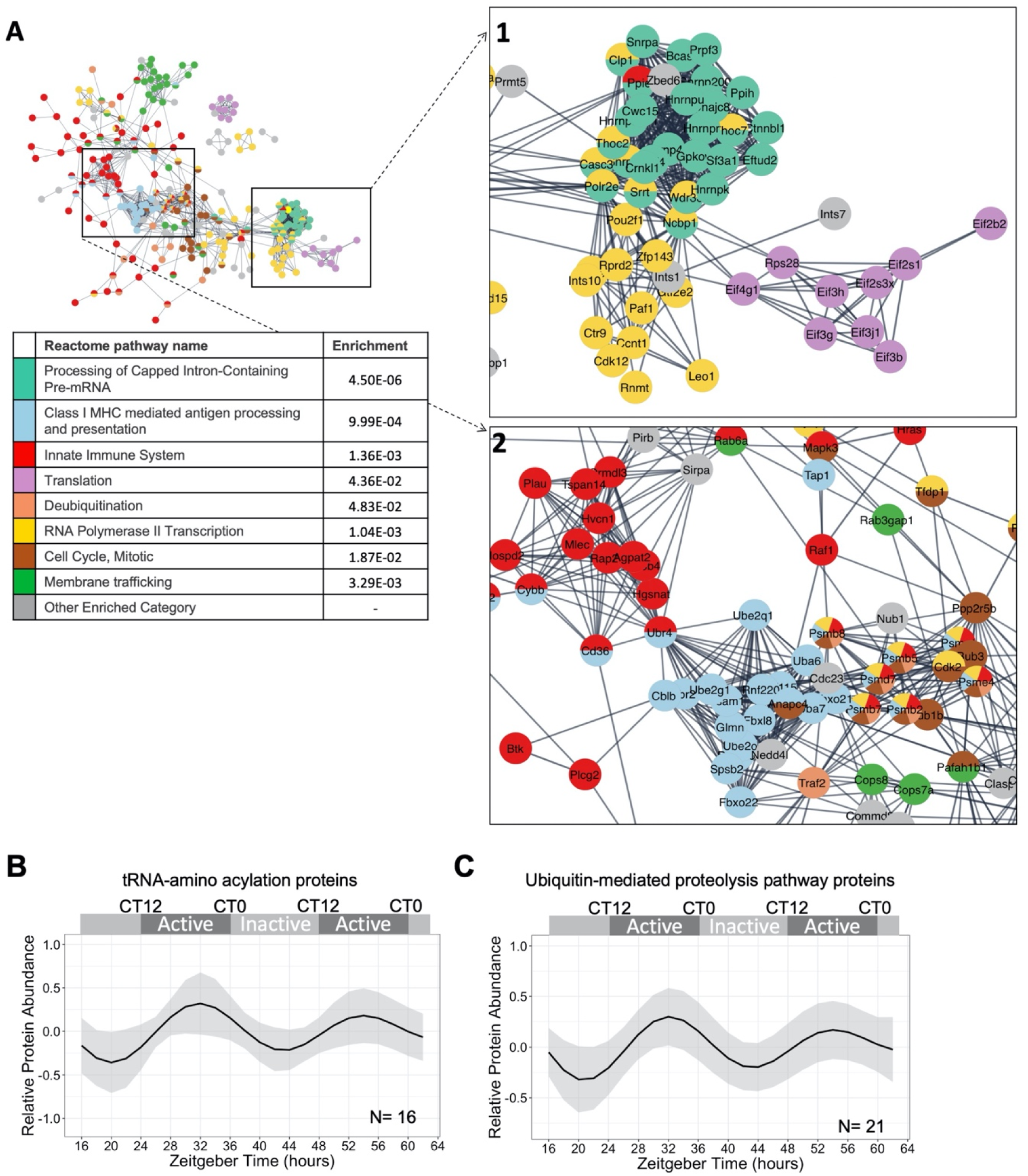
Functional Pathway Enrichments Indicate Late Active Phase Post-Transcriptional/Translational Regulation. (A) A global view of the String-DB network of the proteins in enriched reactome categories that peak in the late active wave (CT15-3), with insets focusing on key interactions, including 1) Class I MHC mediated antigen processing and presentation and Innate Immune System, 2) Translation, RNA Polymerase II Transcription, and Processing of Capped Intron-Containing Pre-mRNA. Interactions shown as strings are filtered to highest confidence interactions (confidence >0.9). Colors denote independent reactome categories. Grey coloring represents genes from enriched reactome categories that did not represent a large proportion of the protein-protein interactome or described redundant categories. (B) An average of the modeled fits of all sixteen circadian proteins in the tRNA-amino acelyation reactome category. Shading indicates +/- 1 standard deviation of models at each time point. (C) An average of the modeled fits of all twenty one circadian proteins in the ubiquitin-proteasome category, as defined by KEGG. Shading indicates +/- 1 standard deviation of models at each time point.

Proteins peaking in the late active wave (CT15-3) were enriched in translational categories (“Processing of Capped Intron-Containing Pre-mRNA”, “RNA polymerase II Transcription”, and “Translation”) and the proteins involved in these processes were highly clustered (Figure 3A, inset 1). Using PSEA, we additionally identified that proteins peaking in the late active phase were involved in tRNA aminoacylation, the ligation of amino acids to tRNA to be used in translation (Supplemental Figure 6A). We then modeled the levels of 16 different rhythmic enzymes catalyzing the formation of tRNA and calculated the average level of tRNA enzymes over the day. We found that the average level of tRNA-synthesizing enzymes peaked during the late active phase (Figure 3B and Supplemental Figure 6B), suggesting that the late active phase is the peak phase for translation. This late active translational phase aligned with the late active peak in protein levels and likely explains the wave of circadian proteins that arise rapidly from circadian RNA at CT18, as well as circadian proteins that arise from non-circadian RNA from CT18-22 (see Figure 2 and Supplemental Figure 3).

However, the late active peak in translational regulation did not explain the peak of circadian proteins near CT10 without associated circadian RNAs. Previous models suggested that a peak in ubiquitin ligases antiphase to protein peak phases would suggest that these proteins are going through timed degradation (Lück *et al.*, 2014). We noted that the late active phase was also enriched for ubiquitin ligases and proteasomal subunits (Figure 3A, inset 2). Moreover, PSEA identified an enrichment for ubiquitin-protein ligases in the late active phase (Supplemental Figure 6A). We modeled and calculated the average level of rhythmic proteins involved in protein ubiquitination/degradation over the day. We found the average peak of protein degrading enzymes was in the late active phase (Figure 3C and Supplemental Figure 6C). This suggests that ubiquitin-targeted proteolysis may explain the peak of circadian proteins near CT10 without associated circadian RNAs.

### Post-transcriptional Circadian Regulation in the Late Inactive Phase Plays an Essential Role in Immunometabolism

With the demonstration of post-transcriptional regulation specific to the late active wave, we next investigated the enrichment of the proteins in the late inactive wave (CT3-15). Highly enriched categories included the “Citric Acid Cycle and Respiratory Electron Transport” and “Metabolism of Vitamins and Cofactors” (Figure 4A). PSEA identified several central energy-producing pathways and mitochondrial cellular components in the late inactive wave (Supplemental Figure 6A). While both the CT15-3 and CT3-15 waves were enriched for “Innate Immunity”, the proteins involved in the late inactive wave (CT3-15) appeared to share more interactions with proteins involved in receptor tyrosine kinase signaling and metabolism (Figure 4A, inset). These data characterize the late inactive wave as a time for energy production via the regulation of central metabolic pathways.

**Figure 4.**
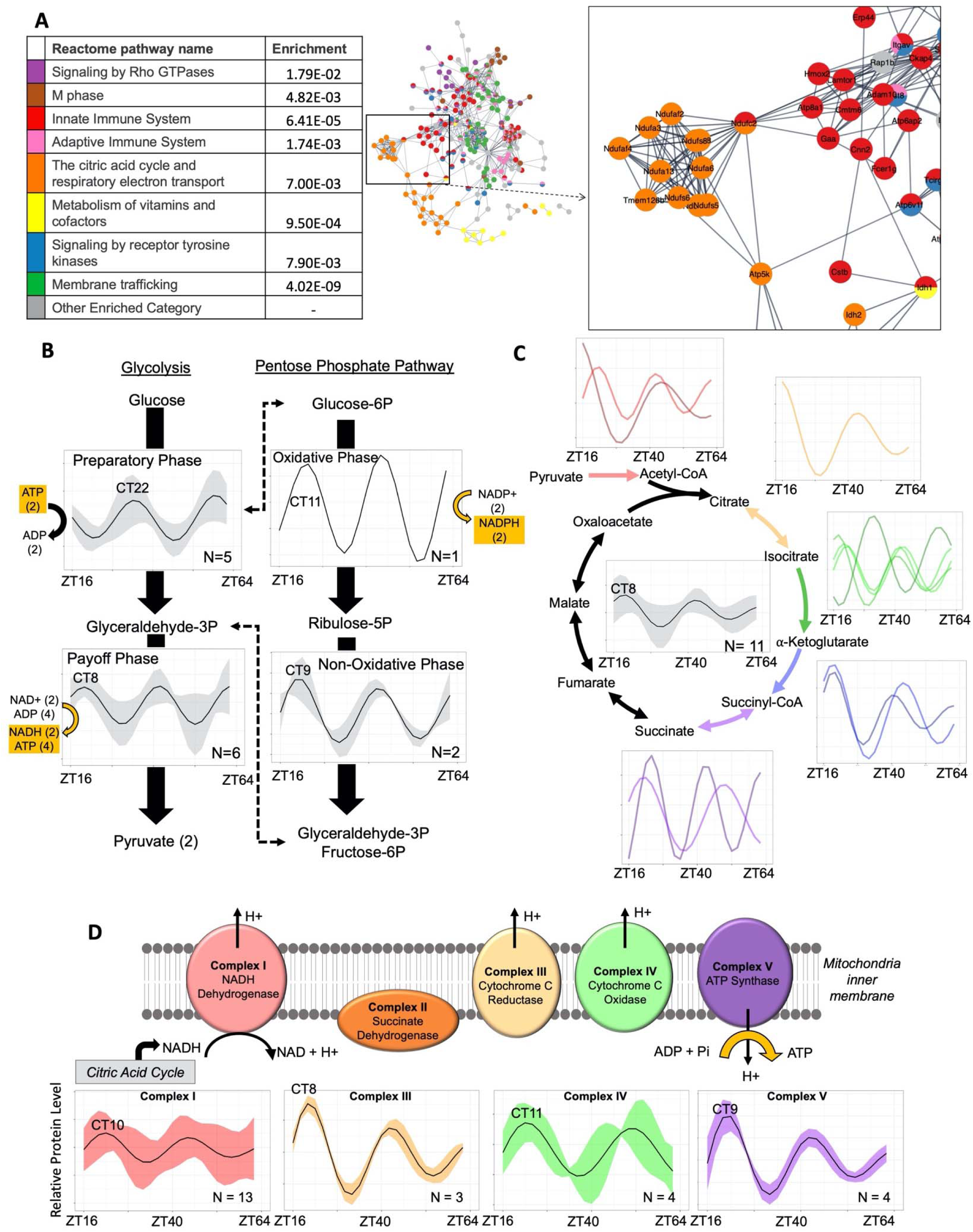
Circadian Regulation of Central Metabolic Pathways Coordinates Mitochondrial ATP Synthesis at Relative Dawn. (A) A global view of the String-DB network of the proteins in enriched reactome categories that peak in the late inactive wave (CT3-15), with an inset focusing on key interactions, including the citric acid cycle and respiratory electron transport. Interactions shown as strings are filtered to highest confidence interactions (confidence > 0.9). Colors denote independent reactome categories. Grey coloring represents genes from enriched reactome categories that did not represent a significant proportion of the protein-protein interactome or described redundant categories. (B) An overview of the oscillations in the payoff/preparatory phases of glycolysis as well as the oxidative and non-oxidative phases of the PPP. The average relative protein abundance model of all circadian proteins identified in each phase is represented, with the shaded region representing the standard deviation between models at each time point. The circadian time (CT) of the first peak is labeled. (C) A schematic of the subunits involved in the TCA cycle which had component proteins identified as rhythmic. An average curve and shaded standard deviation of the overall TCA cycle (center) as well as individual modeled fits for all circadian proteins in the identified corresponding enzyme complex are shown next to each step. The circadian time (CT) of the first peak is labeled. (D) A schematic of the subunits involved in the ETC which had component proteins identified as rhythmic. Average curve and shaded standard deviation of the modeled fits for all circadian proteins in the identified enzyme complex are shown below. The circadian time (CT) of the first peak is labeled.

The timing of the macrophage metabolic state is of interest as metabolic programming is increasingly recognized as a critical factor in determining macrophage immune responses, such as characterizing polarization within the M1 pro-inflammatory/M2 anti-inflammatory spectrum paradigm (Blagih and Jones, 2012; Tang and Mauro, 2017; Van den Bossche, O’Neill and Menon, 2017). To determine if there is circadian regulation of these metabolic pathways in macrophages, we analyzed which proteins circadianly oscillated in the protein complexes involved in the production of ATP (Figure 4B, 4C & 4D). To ultimately produce ATP and other energy mediators (e.g. NADH), glucose is first committed to glycolysis metabolism in the cytosol via the “preparatory” phase. Evaluation of circadian proteins in this critical entrance phase of glycolysis revealed that phosphofructokinases, the rate-limiting first step, as well as enzymatic steps 3 and 4 are circadianly oscillating enzymes that peak during the late active phase (Figure 4B, Supplemental Figure 7A). Conversely, several circadian enzymes representing steps 6, 7, 8 and 9, which is known as the “payoff” phase that results in net ATP/NADH production, peak during the late inactive phase (Figure 4B, Supplemental Figure 7B) (Bonora *et al.*, 2012).

Parallel to glycolysis, the Pentose Phosphate Pathway (PPP) utilizes glucose to produce NADPH. Previous work in *N. crassa* demonstrated circadianly-regulated proteins in the PPP peak in abundance anti-phase to circadianly-regulated glycolysis proteins (Hurley *et al.*, 2018). We found a similar arrangement in macrophages, where the protein PGD from the PPP oxidative phase (NADPH-producing) was circadianly regulated and peaked in abundance in the late active wave, anti-phase to glycolysis preparatory enzymes (Figure 4B, Supplemental Figure 7C). Further, we found that in-phase with glycolysis ATP/NADH-producing phase enzymes, circadianly regulated proteins involved in the non-oxidative phase of the PPP (TKT and TALDO1) peaked (Figure 4B, Supplemental Figure 7C). As the non-oxidative phase of the PPP produces metabolites that can feed back into the glycolysis pathway’s payoff phase, this suggests that energy production via glycolysis vs PPP are primed to enter each pathway at opposite times of the circadian day. Because pro-inflammatory macrophages preferentially use glycolysis to produce metabolites, the in the late active wave suggests that macrophages display a more pro-inflammatory phenotype in the late active wave (Van den Bossche, O’Neill and Menon, 2017).

The product of glycolysis, pyruvate, can be further metabolized in the mitochondria via the tricarboxylic acid (TCA) cycle to generate an abundance of NADH. Eleven protein subunits in five protein complexes in the TCA cycle were circadianly regulated and all but one peaked during the late inactive phase (Figure 4C, Supplemental Figure 7D). Importantly, this included the enzyme catalyzing the initial entry of metabolites derived from glycolysis into the TCA cycle as well as the majority of subunits belonging to protein complexes that facilitate irreversible reactions. Thus, this targeted circadian regulation is likely to propel the TCA cycle in the forward, NADH-producing, direction during the late inactive phase. Moreover, in the electron transport chain (ETC), which utilizes the NADH produced by the TCA cycle to generate large amounts of ATP, twenty-four subunits comprising parts of four ETC complexes oscillated with a circadian period in our dataset (Figure 4D, Supplemental Figure 7E). The average time of proteins reaching their peak levels in the ETC was also in the late inactive phase, slightly delayed from that seen for the TCA cycle. As M2-like, anti-inflammatory macrophages favor ATP production via stable oxidative phosphorylation, our findings suggest that naïve macrophages resemble an anti-inflammatory immunometabolic state in late inactive phase (Van den Bossche, O’Neill and Menon, 2017). To also examine if this circadian regulation of central metabolic pathways is largely derived from post-transcriptional/translational mechanisms, we analyzed which circadian proteins have corresponding circadian mRNA and found that only 4, one in glycolysis, one in pentose phosphate and 2 in the electron transport chain categories were circadian at the mRNA level. Thus, the majority of circadian regulation of metabolic enzymes occurs at the post-transcriptional level in macrophages.

### Post-transcriptional Circadian Regulation Influences ATP Production and Mitochondrial Morphology

Due to the significant circadian post-transcriptional regulation in the TCA cycle and ETC, we hypothesized that there would be a resulting oscillation in the respiratory rate of mitochondria, i.e. macrophage respiratory rate. To analyze oscillations in macrophage respiratory rates, we utilized a Seahorse assay to measure oxygen consumption rate (OCR) at opposing time points over the circadian day (Figure 5A). For each time point, a basal respiratory rate was measured (Figure 5B). As predicted by our analysis of the proteome, where we noted a peak in oxidative phosphorylation protein abundance in the inactive phase, basal respiratory rates were significantly higher during the inactive phase at CT4 compared to CT16 (Figure 5B). OCR was measured after the addition of Oligomycin, an ATP synthase-inhibitor, to provide a measure of ATP production. CT4 exhibited a higher capacity for mitochondrial ATP production as compared to CT16 (Figure 5C). Fluoro-carbonyl cyanide phenylhydrazone (FCCP) was then added to assess the maximal respiration of the mitochondria. We found that maximal respiration decreased as a function of time as opposed to an oscillatory manner (Supplemental Figure 8A). Finally, Rotenone/Antimycin A was added to inhibit mitochondrial function and measure spare respiratory capacity. The spare respiratory capacity decreased with time similar to what was observed for maximal capacity (Supplemental Figure 8B). As a whole, this demonstrates that the circadian clock generates rhythms in basal rate metabolism and ATP-linked oxygen consumption.

**Figure 5.**
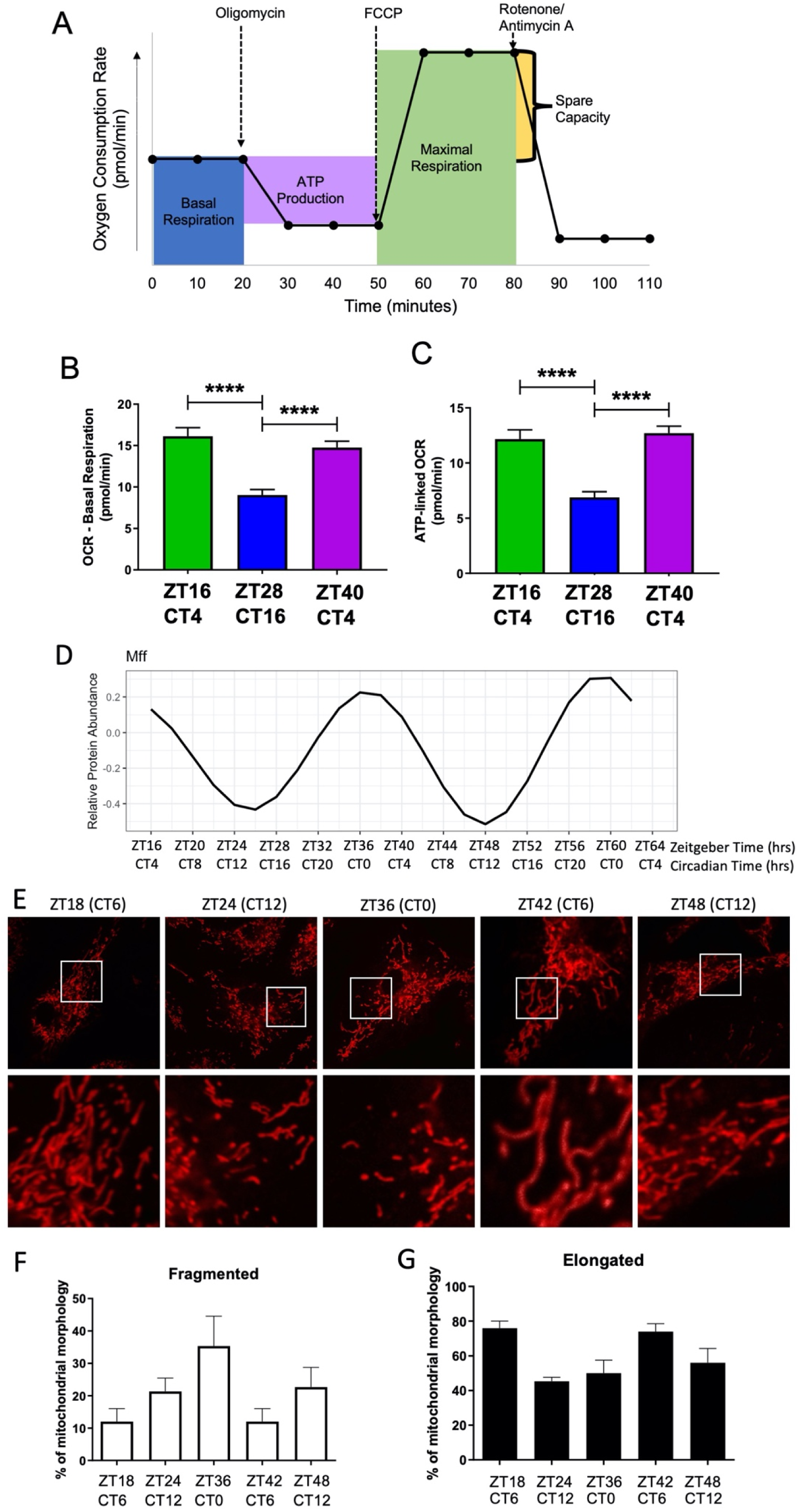
Circadian Regulation of Metabolism and Mitochondrial Morphology Leads to a Peak in Mitochondrial-linked Oxygen Consumption Rate in the Late Inactive Phase. (A) Overview of SEAHORSE protocol showing the frequency of OCR measurement and interpretation of mitochondrial dynamic being measured with each addition of various inhibitors. (B) The overall SEAHORSE traces for BMDMs sampled at ZT16 (CT4), ZT28 (CT16), and ZT40 (CT4) post serum shock. SEAHORSE-derived measurements for the rate of (C) Basal respiration and (D) ATP-linked OCR, in picomoles/minute for 50,000 cells/well and normalized by total DNA per well. (E) The modeled average level oscillation of the MFF protein in the proteomics time course. Across circadian time in synchronized BMDMs, mitochondria were stained using MitoTracker Red CMXRos (F) and the amount of fragmented (G) and elongated (H) mitochondria were quantified by confocal microscopy.

Mitochondria regulate energy production through the coordination of fission (fragmentation of mitochondria) and fusion (elongation of mitochondria). Fused mitochondrial morphology supports higher levels of ATP generation via oxidative phosphorylation (Angajala *et al.*, 2018). Fusion and fission is a dynamic process regulated by a number of proteins. A key protein controlling fission is the Dynamin-related protein 1 (Drp1), a GTPase which is recruited to the mitochondrial outer membrane by Mitochondrial fission factor (Mff) (Smirnova *et al.*, 2001; Otera *et al.*, 2010). In our dataset, MFF protein oscillated in a circadian manner, but its transcript did not (Figure 5D). When we surveyed additional proteins involved in the fission/fusion pathway, we found that Fis1 was also rhythmic (Supplemental Figure 8C, Supplemental Table 2). We postulated that the oscillation of Mff would lead to an oscillation in mitochondrial morphology that would also coincide with the observed oscillations in basal/ATP-linked oxygen consumption. To investigate this, mitochondrial morphology was assessed over time by imaging macrophages stained with MitoTracker Red (Figure 5E). Indeed, a peak in the percentage of fragmented, i.e. fissioned, mitochondria was observed in the late active phase at CT0/24 (Figure 5F), coinciding with the peak in MFF protein abundance (Figure 5D). The highest percentage of elongated, i.e. fused mitochondria occurred at CT6, coinciding with the time of day with highest basal and ATP-linked oxygen consumption (Figure 5G).

### Circadian Regulation of Metabolism Impacts the Phagocytic Immune Response

Mitochondrial function is highly influential to immune responses (Mills and O’Neill, 2016). We thus proposed that circadian regulation of mitochondrial dynamics and ATP production would have distinct consequences on a macrophage’s response to immunological stimuli. One of the characteristic functions of macrophages in their role as mediators of the immune response and inflammation is their phagocytic activity. Phagocytosis involves complex networking of surface receptors, signaling cascades, and cytoskeletal modifications, making phagocytosis an energy-demanding process requiring interconnection with cellular metabolism. In addition to metabolic regulation, PSEA for circadian proteins identified in the active wave the ontological category “Positive Regulation of I-Kappa B Kinase NF-kB Cascade”, a category that highlights proteins involved in the signaling pathway that leads to the phosphorylation and degradation of I-Kappa B, enabling a robust inflammatory response via NF-kB signaling (Figure 6B, Supplemental Figure 3C) (Liou, 2002).

**Figure 6.**
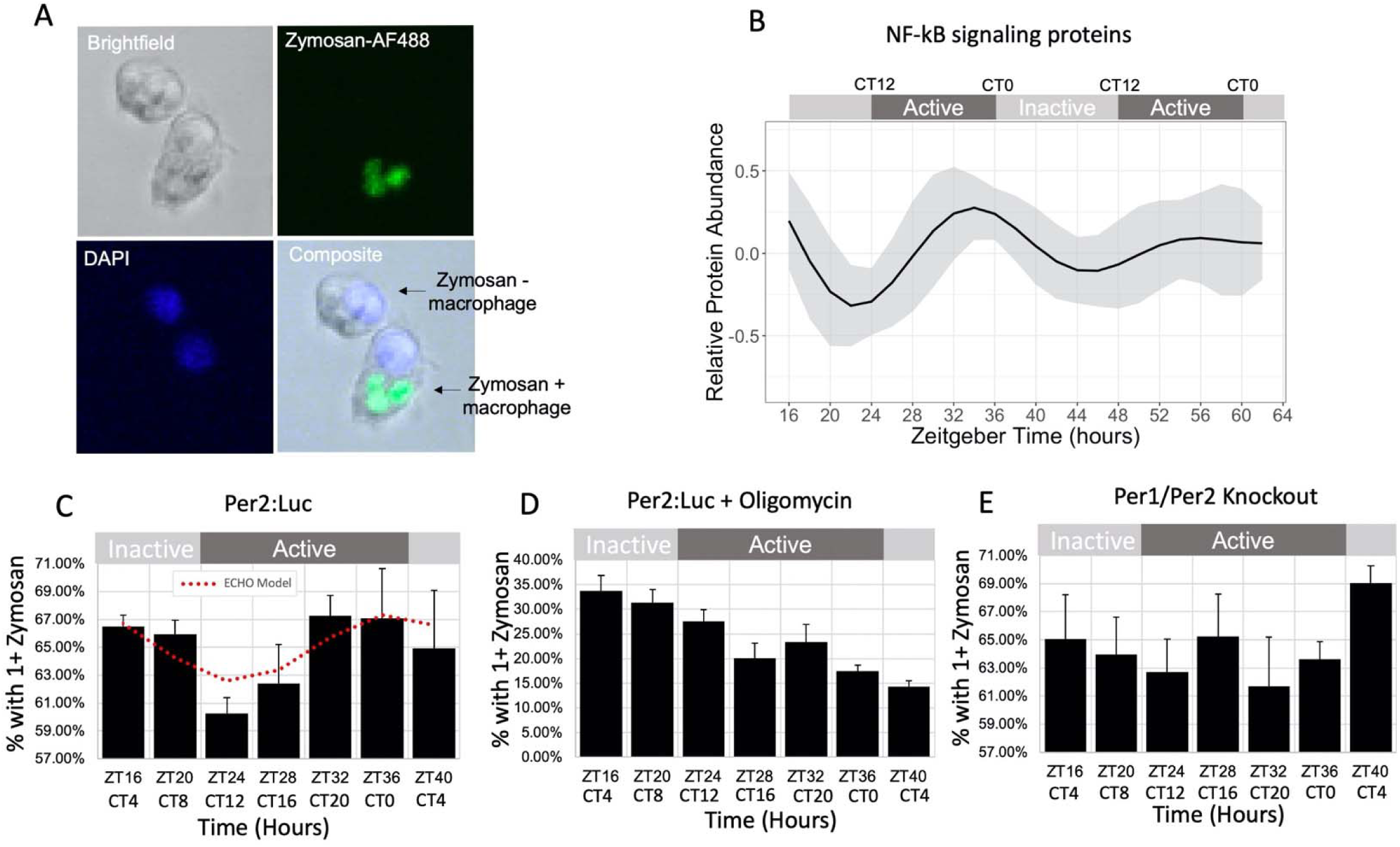
Macrophage Phagocytosis of Zymosan Shows Metabolism-Related Circadian Variation *in vitro*. (A) Representative confocal microscopy images demonstrating identification of macrophages positive and negative for Zymosan-AlexaFluor488 phagocytosis. Cells were also DAPI stained to aid in cell counting. (B) An average of the modeled fits of all sixteen circadian proteins in the NF-kB signaling pathway. The levels of phagocytosis is reported as a percent of cells with one or more particles of Zymosan in (C) Per2::Luc, (D) Per2::Luc co-incubated with Oligomycin, and (D) Per1/Per2 knockout BMDMs at a 4hr resolution for 24 hours. Significance trace of oscillation as determined by ECHO is plotted on the Per2::Luc graph in dotted red lines. Error bars represent Standard Error of the Mean (SEM).

To determine if macrophage phagocytic activity was circadianly timed in our *in vitro* model, we measured the phagocytosis of AF488-conjugated Zymosan, a yeast-derived bioparticle, as Zymosan had previously been shown to be rhythmically phagocytized by ex-vivo murine peritoneal macrophages and is also known to exert inflammatory signaling primarily via TLR2/Dectin-1 receptors, which have cooperating signaling cascades converging on NF-kB signaling (Figure 6A) (Gantner *et al.*, 2003; Dennehy and Brown, 2007; Hayashi, Shimba and Tezuka, 2007). Luminescence tracking further examined that the addition of zymosan and/or oligomycin was not radically altering clock gene expression for the duration of our experiment or ablating Per2 rhythmicity, though long term, immune challenge at CT4 decreased the amplitude of the oscillation whereas immune challenge at CT16 did not. (Supplemental Figures 9A - C). Consistent with previous studies of on circadian phagocytosis, BMDMs derived from Per2:Luc mice were observed to rhythmically phagocytize Zymosan, with the lowest phagocytosis rate at CT12 (late inactive phase) and the highest at CT0 (late active phase) (Figure 6C), with an 11.4% increase in efficiency at the peak vs the trough (Hayashi, Shimba and Tezuka, 2007; Oliva-Ramírez *et al.*, 2014).

Both ECHO and JTK analysis confirmed the significance of this rhythmicity (Supplemental Table 3) (Hughes, Hogenesch and Kornacker, 2010; De los Santos *et al.*, 2017, 2020). To determine if circadian metabolic regulation was the driving force in the rhythmic regulation of phagocytosis, we added the ATP synthase inhibitor Oligomycin A to the phagocytosis assay at each timepoint and measured phagocytosis 30 minutes later (Figure 6D) (Jastroch *et al.*, 2010). We noted a dramatic decrease in phagocytosis that was exacerbated over time, with no evidence of oscillation of phagocytosis in these cells (Figure 6D). To eliminate the possibility that the phagocytosis rhythm was dictated independently of clock function, we analyzed phagocytosis rates from BMDMs derived from Per1/Per2 double knockout mice, which are arrhythmic in all tissues (Bae *et al.*, 2001; Zheng *et al.*, 2001) and found that phagocytosis was not rhythmic in the Per1/Per2 knockout BMDMs (Figure 6E). Modeling the levels of the signaling proteins involved in the I-KappaB Kinase/Nf-kB cascade demonstrated that these proteins peaked at the same time as phagocytosis (Figure 6B, 6C and Supplemental Figure 9D), suggesting that circadian signaling pathway regulation in crosstalk with circadian metabolic regulation could also play a role in the generation of the rhythms observed.

## DISCUSSION

Our comprehensive profiling of the macrophage transcriptome and proteome over circadian time revealed extensive circadian post-transcriptional regulation (Figure 1 and Supplementarpy Figure 2) with two major transcriptional/translational waves occurring anti-phase in either the late-active or -inactive phases (Figure 2). Our previous study in the fungal clock model *Neurospora crassa*, and others’ less-powered studies in other mammalian tissue types, have suggested as much as 40% of the circadian proteome is post-transcriptionally regulated (Reddy *et al.*, 2006; Kojima, Shingle and Green, 2011; Chiang *et al.*, 2014; Hurley *et al.*, 2014, 2018; Mauvoisin *et al.*, 2014; Robles, Cox and Mann, 2014; J. Wang *et al.*, 2017; Green, 2018; Wang *et al.*, 2018; Mauvoisin and Gachon, 2019). Our study observed a marked increase in the lack of overlap between circadian mRNA and protein as compared to these previous studies. Of the 1,500+ circadian proteins we identified, 85% of those proteins did not have corresponding rhythms in mRNA expression and only 32% of circadian mRNA resulted in a rhythmic protein. In addition, there were far more circadian proteins than circadian mRNAs of the subset of genes detected in both datasets. This indicates that post-transcriptional/translational mechanisms play a particularly significant role in macrophages, so much so as to be a challenge to the canonical view that positive-arm transcriptional regulation is the dominant driver of circadian output in all tissues. This discordance may extend to all mammalian tissue types, as previous studies have not been able to achieve the power of our extended, densely sampled, study (Reddy *et al.*, 2006; Kojima, Shingle and Green, 2011; Hurley *et al.*, 2014, 2018; Mauvoisin *et al.*, 2014; Robles, Cox and Mann, 2014; Green, 2018; Wang *et al.*, 2018; Mauvoisin and Gachon, 2019). It is possible, however, that macrophages are unique in the extent of their post-transcriptional regulation. This regulation may be necessary due to the macrophages’ need to respond quickly to a wide variety of potential immunological stimuli (Abbas, Lichtman and Pillai, 2018). Future work deeply evaluating proteomes from other tissues, as well as the generation of a complete network of the circadian transcriptional cascade, is needed to determine if this is the case.

Post-transcriptional regulation in mammalian tissues has been suggested to primarily derive from systemic cues (Koike *et al.*, 2012; Kojima, Sher-Chen and Green, 2012; Menet *et al.*, 2012; Partch, Green and Takahashi, 2014). However, our data shows that significant post-transcriptional regulation occurred in macrophages devoid of systemic cues *in vitro*, contradicting this hypothesis. In contrast to solid tissues, where stochasticity between millions of cells can be buffered against in producing an overall circadian output from a tissue, macrophages are largely independent in their surveillance and initiation of early stages of inflammation (Nagoshi *et al.*, 2004; Forger and Peskin, 2005; Lande-Diner *et al.*, 2015). This suggests that their intrinsic clock output would need to be a finely-tuned, powerful driver of advantageous rhythms, thus utilizing post-transcriptional regulation to create oscillations in protein abundance may be an essential circadian regulatory mechanism unique to macrophages.

In addition, differing from *N. crassa* and liver cells, it appears that degradation and transcription mechanisms both may play a large role in macrophage circadian regulation, suggesting that different tissue types in the mammalian clock, and perhaps specifically the macrophage clock, have evolved more complex mechanisms of post-transcriptional regulation (Figure 3) (Lück *et al.*, 2014; Caster *et al.*, 2016; Hurley *et al.*, 2018). In addition to giving rise to rhythmic proteins without rhythmic mRNA, the robust circadian control of protein degradation and transcription we noted (Figures 2 and 3) may also give rise to rhythmic expression of transcripts that do not produce rhythmic proteins. Ribosome and degradation profiling would shed light on these relationships, as this cannot be deciphered by our steady state measurements. Our findings also highlight the need for circadian studies that utilize multi-omics approaches, as transcription-focused interrogations are largely inadequate for understanding physiologically relevant regulation stemming from the circadian clock. Further, the difference in rhythmic transcripts between tissue types, coupled with differences in post-transcriptional regulation, suggest that a great deal more physiology could be learned from studies across multiple tissue types (Figure 2) (Hurley *et al.*, 2018; Mure *et al.*, 2018).

Our deeply sampled dataset served as a rich resource to investigate the impact of circadian regulation on macrophage physiology, demonstrating that the circadian clock generates distinct immunometabolic states at opposing times of day. During the inactive phase, macrophages were especially primed for ATP synthesis via oxidative phosphorylation as they possessed high proportions of elongated mitochondria and metabolic enzymes for the TCA cycle and electron transport chain were most abundant (Figures 4 & 5). Oxygen consumption rate measurements confirmed that these oscillations imparted an increase in the basal metabolic rate during the inactive phase, though surprisingly not in the maximal capacity. However, this lack in oscillation of maximal capacity may simply be due to the technical limits of the assay. As the circadian day progressed into the active phase, proteins for committal into the glycolysis pathway peaked in abundance and the percentage of fissioned mitochondria reached its peak, with a decreased oxygen consumption rate (Figures 4 & 5). Also, during this time zymosan was most readily phagocytized, suggesting that these distinct immunometabolic states impact immunological responses (Figure 6C). In further evidence for this interconnection, arrhythmic macrophages showed no phagocytic rhythms (Figure 6E), yet similar average levels of phagocytosis, indicating that intrinsic circadian clock output is manipulating these rhythms. Finally, macrophages co-incubated with mitochondrial ATP synthase inhibitor showed no phagocytic rhythms and greatly reduced the amount of phagocytosis, showing that ETC function plays an active role in phagocytosis and is a particularly energy-sensitive process (Figure 6D).

In summary, our findings reveal a paradigm where post-transcriptional circadian regulation times naïve macrophages during the inactive, rest phase to be metabolically primed to resemble an anti-inflammatory phenotype (Figure 7), which characteristically has intact, robust ATP production via mitochondrial oxidative phosphorylation. Upon entering the active phase, the immunometabolic state resembles a pro-inflammatory programming, as reliance on glycolysis over oxidative phosphorylation is a hallmark of pro-inflammatory macrophages (Van den Bossche, O’Neill and Menon, 2017; Carroll *et al.*, 2019). A circadianly-timed proclivity for inflammatory responses in the active phase was consistent with our findings that zymosan is more readily phagocytized at the end of the active phase (Figure 6C) as well as the majority of studies in both humans and nocturnal animals, which show that immune challenges are generally less effectively neutralized when exposed during the inactive phase or under circumstances of circadian disruption when macrophages are not primed for appropriate inflammatory responses (Geiger, Fagundes and Siegel, 2015; Nagy and Haschemi, 2015; Van den Bossche, O’Neill and Menon, 2017). Evolutionarily, it is logical that the circadian clock would advantageously prepare macrophages to respond to immunological threats during the active phase when the majority of pathogenic exposure would occur and then during the inactive phase support a phenotype associated with tissue repair when pathogenic threats are lower. As circadian control of metabolic pathways has been shown to be conserved in a multitude of tissue types and organisms, our study serves as a rich foundation to generate hypotheses about the diverse functional impacts of metabolic regulation in a multitude of biological and disease contexts.

**Figure 7.**
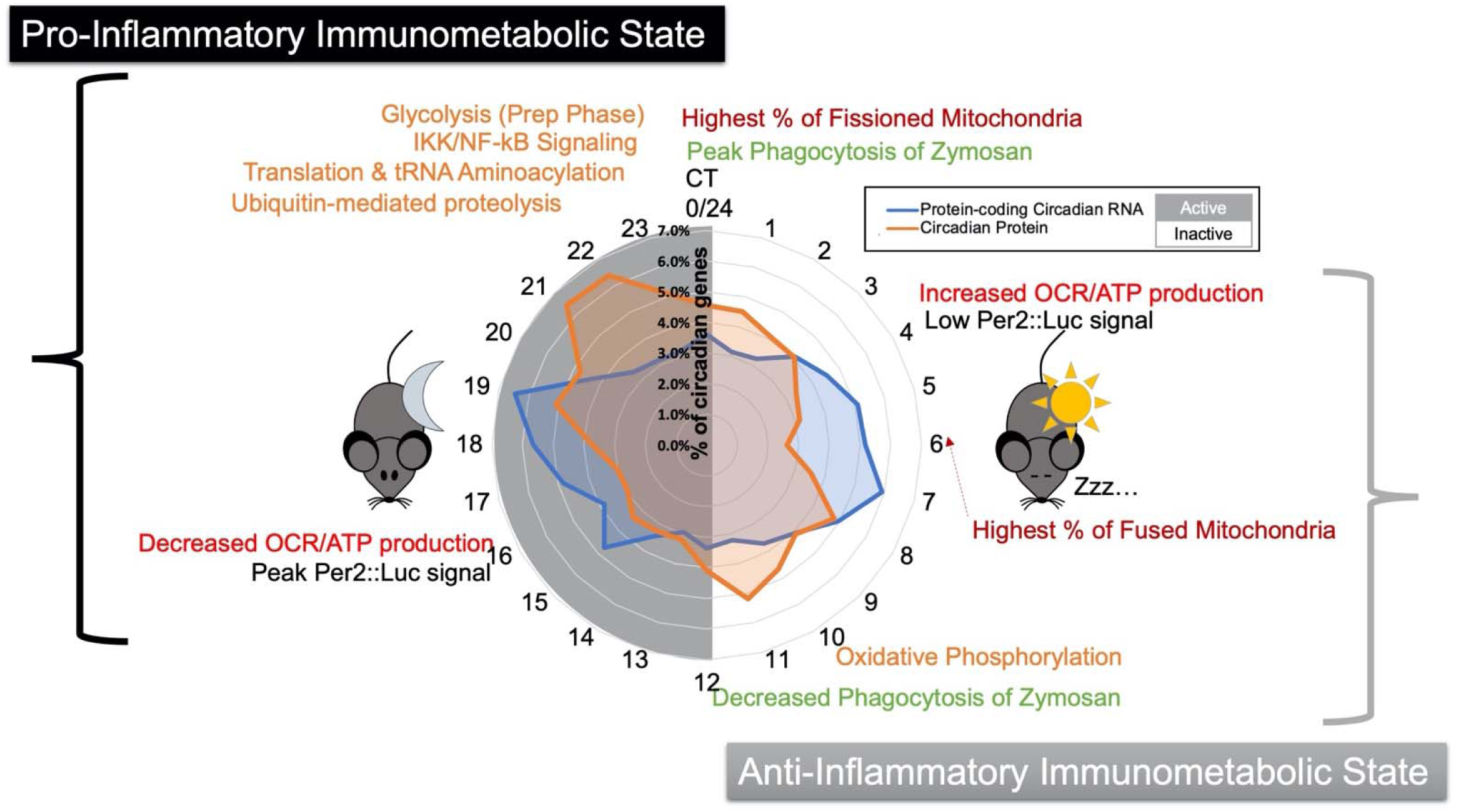
Summary Model of the Circadian Regulation of Distinct Immunometabolic States. A summary schematic displaying key findings spatially organized by circadian time of day, with a radial histogram in the center to indicate when circadian proteins and circadian mRNAs peak in abundance. Orange text represents enriched categories identified in the proteome, red represents the occurrence of mitochondrial morphology/functional observations, green represents phagocytosis assay observations and black text shows the trough and peak of PER2 levels as determined by Per2::Luc luminescence signal. For reference, ZT16 is equivalent to CT4.

As chronic circadian disruption via shift work, increased exposure to light at night, and inadequate daylight exposure become increasingly pervasive in modern society, so have we seen an increase in diseases linked to chronic circadian disruption, such as cancer, obesity, diabetes and cardiovascular and neurodegenerative disease (Evans and Davidson, 2013; Musiek and Holtzman, 2016; Shostak, 2017). Inflammation plays an important part in the pathology of all these diseases, leading many to hypothesize that the dysregulation of immunity plays a major role in the link between chronic circadian disruption and associated diseases. Our work provides profound mechanistic evidence for this link as disruption to the intricate metabolic and clock regulatory networks we identified in macrophages could result in a dysregulation of immunity, potentially fueling chronic inflammation that contributes to disease. Our datasets and analyses provide a robust foundation on which we can build to understand the role of circadian clocks in macrophages to ultimately inform novel strategies for prevention and treatment of diseases associated with chronic circadian disruption and inflammation.

## METHODS

### Animals

PER2::LUC C56BL/7 mice were acquired from Jackson Labs (Accession # 006852) and housed under 12:12 LD conditions with ad libitum access to standard rodent chow. Male mice between the ages of 3-6 months were used for all omics and phagocytosis experiments and were euthanized for tissue collection via CO^2^ inhalation/cervical dislocation.

### Time Course Cell Synchronization

For each time course, bone marrow was harvested from femurs and tibias of 1 mouse with needle and syringe and red blood cells were lysed (Zhang, Goncalves and Mosser, 2008). Cells were then plated at a density of 1*10^6^ in DMEM supplemented with recombinant M-CSF, 10% Fetal Bovine Serum (FBS), L-glutamine, penicillin/streptomicin, and gentamicin. After 3 days incubation at 37°C in 5% CO^2^, additional fresh media was added and incubated another 3 days. Non-adherent cells were then washed off with warmed PBS and media replaced with Leibovitz supplemented with MCSF and 10% FBS for 24 hours. Plates were again washed with warmed PBS and cells starved for 24 hours in Leibovitz with MCSF. Cells were then serum shocked for 2 hours in Liebovitz with 40% FBS and MCSF and then washed with PBS and assayed in Liebovitz with 10% FBS, MCSF and Luciferin (Balsalobre, Damiola and Schibler, 1998). Plates were sealed with grease and glass cover slips. Luminescence over time was recorded using a LumiCycle32 luminometer (Actimetrics) to confirm synchronization of circadian rhythmicity.

### Flow Cytometry

To confirm uniform differentiation of bone marrow cells into macrophages, cells were probed for macrophage-specific markers CD11b and F4/80 both after 6 days of DMEM incubation as well as at the completion of each time course. Cells were lifted off the plate with Cell Stripper, washed with PBS, and incubated with Fc-receptor blocking antibodies to prevent non-specific binding and Zombie Violet to determine proportions of viable and dead cells. Heat killed cells were included as a Zombie Violet positive control. Cells were then incubated with anti-CD11b(AF488) and anti-F4/80(PerCP/Cy5.5) fluorescently conjugated antibodies (Invitrogen) and assayed with a BD LSRII flow cytometer.

### Proteomics

Every 2 hours over 48 hours post-serum shock, protein was extracted to determine global proteome changes over 2 circadian cycles. Triplicate independent time courses were completed for a total of 72 samples. At each time point, cells were lysed with 8M Urea buffer (8.5 pH) and lysate was flash frozen in liquid nitrogen. Multiplex proteomics was performed by Harvard’s Thermo Fisher Center for Multiplexed Proteomics using Tandem Mass Tag Mass Spectrometry (Supplemental File 2-will be published upon peer review). Protein Quantification was performed using the micro-BCA assay by Pierce and after quantification samples were reduced with DTT, alkylated, and an ∼200µg aliquot was precipitated by Methanol Chloroform. Proteins were resuspended in 6M urea and digested overnight with LysC (1:50 protease:protein ratio) and Trypsin (1:100 protease:protein ratio). 8 total multiplexes were run with each containing 50 ug each of protein from 9 randomized time point samples and one common pooled sample between all multiplexes. One complete set of 12 fractions from the total proteome Basic pH reverse phase fractionation from all experiments was analyzed on an Orbitrap Lumos mass spectrometer. Peptides were identified by their MS2 spectra and mapped to corresponding proteins using the SEQUEST algorithm with UniProt reference *Mus musculus* proteome UP000000589. Peptide spectral matches were filtered to a 1% false discovery rate and proteins were quantified only from peptides with a summed SN threshold of >=200 and MS2 isolation specificity of 0.5.

### RNA-seq

Every 2 hours over 48 hours post-serum shock, RNA was extracted to determine global transcriptome changes over 2 circadian cycles. Triplicate independent time courses were completed for a total of 72 samples. At each time point, cells were lysed with Qiagen’s Buffer RLT and lysate was homogenized with Qiashredder columns and flash frozen in liquid nitrogen. All samples were then processed on the same day with Qiagen’s RNeasy kit with on-column DNase digestion. rRNA depletion and RNA-seq was completed by Genewiz on an Illumina HiSeq configured with 2×150 bp paired-end output. Sequence reads were trimmed for adapter sequences then mapped to *Mus musculus* GRCm38.87 reference genome using DRAGEN alignment software. Hit counts were calculated with HTseq and then normalized into transcripts per million (TPM) format.

### Data Processing and Detection of Rhythms

RNA-seq and proteomics data were further processed to remove batch effects using the LIMBR pipeline with default parameters and with removal of peptides with more than 30% missing data points followed by imputation of missing data, as required by LIMBR (Crowell *et al.*, 2019). Batch effect-adjusted data was then analyzed for rhythms with ECHO in free-run mode with smoothing and normalization options (De los Santos *et al.*, 2020). Genes were considered circadianly oscillating if BH-adjusted p-value <.05, forcing coefficient between 0.15 and -0.15, as recommended in (De los Santos *et al.*, 2020) as well as if the period was between 20-28 for RNA and 18-30 for proteins, based on evaluating the known clock gene periods as modeled by our ECHO analysis method. To compare the circadian times at which transcripts/proteins peak, CT phase times were adjusted by period length. To compare genes between datasets, uniprot numbers were used to look up current MGI primary gene names. Duplicate circadian proteins due to database differences or multiple isoforms present were manually curated by choosing a representative protein based on which had more peptides quantified.

### Reactome Enrichment & String-DB Analysis

List of circadian proteins generated with ECHO was bisected into two groups based on circadian adjusted time of peak: those peaking between CT3-15 and those peaking between CT15-3. Each list was analyzed with PantherDB’s PANTHER overrepresentation test using Reactome version 65 (2019-03-02) against the full *Mus musculus* database. Categories were considered enriched if the Bonferroni corrected p-value was <.05. For String-DB visualization and analysis, both lists were subsetted to only include proteins that belong to at least one enriched category. String-DB networks were then generated and colored by Reactome category using the stringApp for Cytoscape and selecting to visualize only high confidence interactions (>.9) (Doncheva *et al.*, 2019).

### Phase Set Enrichment Analysis

Phase set enrichment analysis for circadian proteins was completed using the downloaded .jar tool and with default analysis parameters as defined by (Zhang *et al.*, 2016). Murine gene ontology gene sets were downloaded from Gene Set Knowledgebase (Bares and Ge, 2019). Enrichment was determined versus a uniform background and using the standard cutoff of adjusted p<.05.

### Zymosan Phagocytosis Assay

Macrophages were differentiated/grown to confluence, serum shocked as described above utilizing two 12-hour staggered groups and assayed in a 32-plate lumicycle (Actimetrics). Starting at ZT16, every 4 hours over 24 hours, in triplicate, 11 ul of 20 mg/ml zymosan per plate (∼1:100 cell:zymosan ratio) was added directly to assay media with macrophages and returned to lumicycle for incubation at 37C for 30 minutes. Prior to incubation, AF488 labeled Zymosan (Invitrogen, cat #: Z23373) was opsonized with FBS from the same lot as used for macrophage media according to manufacturer’s instructions. For oligomycin time course, oligomycin was added to the media at the same time as zymosan to a final concentration of 1.5uM. After incubation, cells were washed 3X with PBS and lifted from plates using CellStripper. Cells were then fixed in Neutral Buffered Formalin for 30 minutes in the dark at room temperature. Cells were then resuspended in PBS and stored at 4C until imaging. On day of imaging cells were stained with DAPI and imaged with a Zeiss LSM 510 laser scanning confocal microscope using a 40X objective until well over 100 cells were imaged for each sample. For each image, first the number of total macrophages was counted in ImageJ using “CellCounter” plugin with the composite of brightfield/DAPI channels. Only macrophages with clearly defined borders entirely within the image bounds and a DAPI stained nucleus were considered suitable for analysis. Then the number of macrophages with positive Zymosan-AF488 signal was counted to calculate of the percent of zymosan-positive macrophages out of at least 100 total macrophages observed per time point.

### Mice for Mitochondrial Experiments

WT Mice used for experiments were both male and female, aged between 8-12 weeks. These mice were bred and maintained in specific pathogen free conditions in the comparative medicine unit, Trinity College Dublin. All mice were maintained according to European Union regulations and the Irish Health Products Regulatory Authority.

### Bone Marrow Derived Macrophage Differentiation for Mitochondrial Experiments

Bone marrow derived macrophages (BMDMs) were prepared from the femurs and tibia of mice. Mice were sacrificed by carbon dioxide inhalation and cervical dislocation. Hair, skin, and muscle was removed from each bone and both ends were cut using a scissors. A syringe was used to flush bone marrow with warm complete Dulbecco’s Modified Eagle’s Media (cDMEM; 10% fetal bovine serum, 1% penicillin/streptomycin) which was collected into a Falcon tube. The cell suspension was centrifuged, the supernatant was discarded, the cell pellet was resuspended in red blood cell lysis buffer, and after 3 minutes warm cDMEM was added to deactivate the lysis buffer. The cell suspension was passed through a 40μm mesh filter into a Falcon tube, centrifuged, and the supernatant was discarded. The cell pellet was resuspended in warm cDMEM supplemented with 10% L-929 fibroblast supernatant. The cell suspension was divided between 3 10cm cell culture plates and incubated in a humidified 5% CO2 incubator at 37°C. After 6 days of growth, cells were scraped in ice cold Dulbecco’s phosphate buffered saline (PBS), collected in a falcon tube, and centrifuged. The supernatant was discarded and the cell pellet was resuspended in cDMEM. Viable macrophages were counted by haemocytometric analysis using the Countess Automated Cell Counter (Invitrogen) according to manufacturer protocol and equalized to an appropriate density.

### Seahorse

50,000 BMDMs were seeded in each well of a 96-well Seahorse plate. Cells were left to adhere for several hours in a humidified 5% CO2 incubator at 37°C in cDMEM supplemented with 10% L-929 medium. Cells were serum shocked to T16, T28, or T40 and the start time of the serum shock for each group was staggered so that each time could be analysed by Seahorse assay simultaneously. At the appropriate time to begin the serum shock protocol for each group, media was changed to complete Leibovitz L-15 media supplemented with 10% L-929 media and 1% HEPES for 12 hours. Cell media was then changed to serum free DMEM supplemented with 1% P/S and 10% L-929 media for 12 hours. Cells were then serum shocked by replacing cell media with DMEM supplemented with 50% horse serum and 1% P/S for 2 hours. Following this, media was replaced with cDMEM supplemented with 10% L-929 medium and cells were incubated for an appropriate time before beginning the Seahorse assay. Following completion of the serum shock protocol for all groups, cells were washed with Seahorse XF DMEM Medium pH 7.4 (Agilent, 103575-100) supplemented with 10mM glucose, 1mM sodium pyruvate, and 2mM glutamine. Cells were then incubated in a non-CO2 incubator at 37°C for 45 minutes. Mitochondrial stress test compounds (1.5mM oligomycin, 1mM FCCP, 0.5mM rotenone/antimycin a) were prepared in supplemented Seahorse XF DMEM Medium and loaded into the appropriate ports of an XFe96 Seahorse sensor cartridge. Following calibration, cells were loaded into the Seahorse XFe96 instrument and a mitochondrial stress test was carried out. Following completion of the assay normalization was carried out via PicoGreen dsDNA quantitation.

### Mitochondrial Dynamics

BMDMs (0.5 × 10^6^) were plated on µ-Dish 35 mm, high Glass Bottom (Ibidi, Germany) and maintained overnight at 37 °C in a 5% CO_2_ atmosphere. The cells were synchronized by serum shock as described by (Balsalobre, Damiola and Schibler, 1998). After serum shock, cells suspended in fresh 5% FBS Dulbecco’s Modified Eagle Medium. Thirty minutes before the indicated time, cells were loaded with MitoTracker Red CMXRos (Life technologies) at a final concentration of 50 nM. Cells were washed with PBS fresh; medium was placed, and cells were observed in a Leica SP8 scanning Confocal (Wetzlar, Germany), with a 63x immersion objective. Cell images were analyzed with the ImageJ software (National Institutes of Health, Bethesda, MD). For ImageJ analysis, the mitochondrial length of more than 50 mitochondrial particles per cell was measured in over 25 cells per experimental condition, out of three independent experiments. Mitochondria were then divided into two different categories, based on length, as mitochondria of less than 1 mm (fragmented) and greater than 3 mm (elongated), as described by (Park et al., 2013).

## ACKNOWLEDGEMENTS

We would like to thank Dr. Kristen Bennett and Hannah De los Santos of the Rensselaer Institute for Data Exploration and Applications for their help with ECHO analyses. Gretchen Clark at Rensselaer for her help with macrophage work and other members of the Hurley lab for their valuable discussion and feedback. Sergey Pryshchep of the Rensselaer Microscopy Core for his technical expertise and the Bioresearch core for animal husbandry. The Dunlap lab at Dartmouth for generously sharing Per1/Per2 knockout mice. This work was supported by the NIH National Institute of General Medical Sciences (T32GM067545 to E.J.C. and GM128687 to J.M.H.); Rensselaer startup funds (to J.M.H.); Consejo Nacional de Ciencia y Tecnología (CVU440823 to M.P.S); Science Foundation Ireland Career Development Award (17/CDA/4688 to A.M.C.).

