## Supplemental Figures for "Post-Transcriptional Circadian Regulation in Macrophages Organizes Temporally Distinct Immunometabolic States"

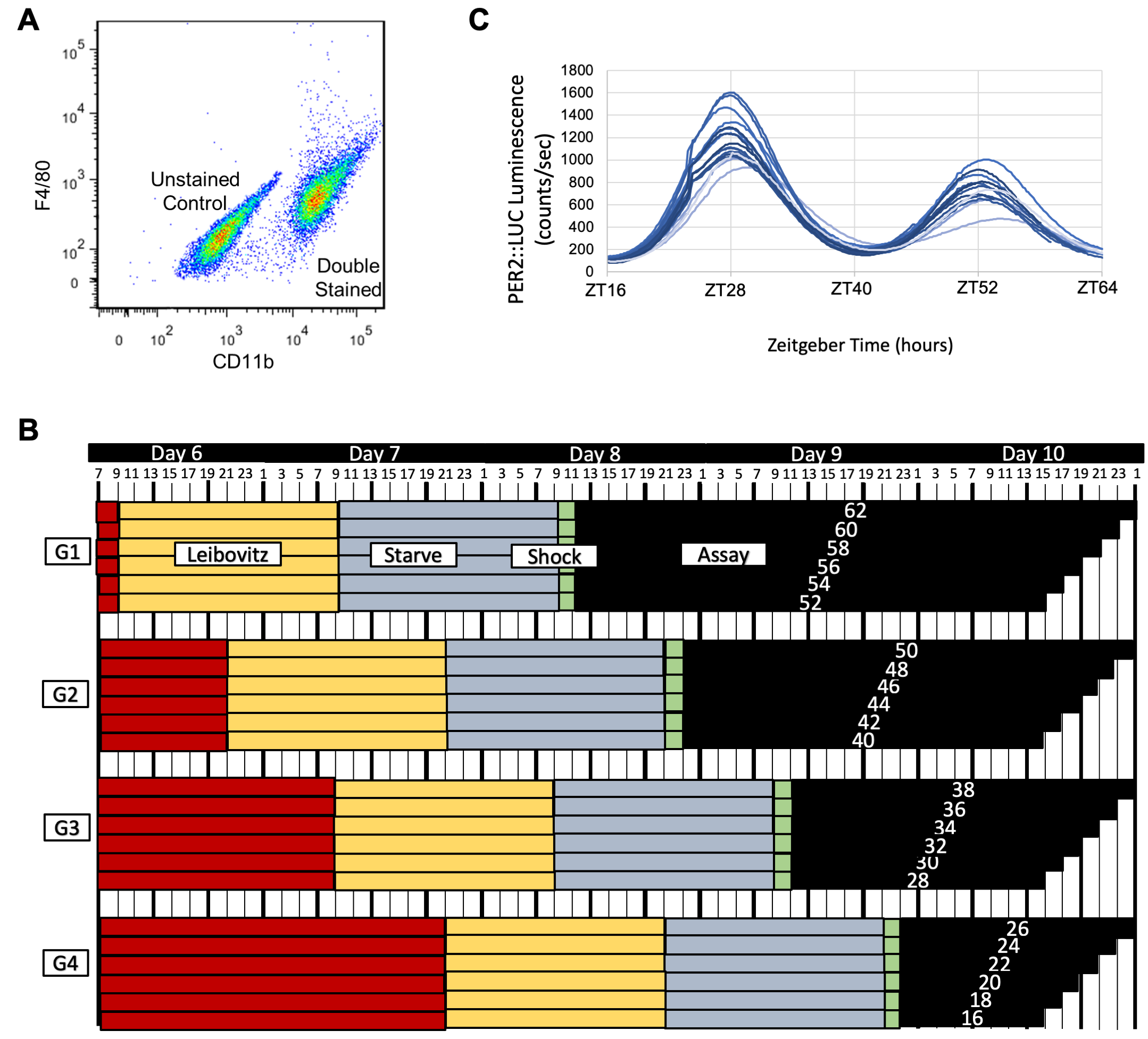

**Supplemental Figure 1. BMDMs Provide Robust Circadian Time Courses.**

(A) A Representative flow cytometry plot confirming the near total differentiation of bone marrow stem cells into macrophages using markers for F4/80 and CD11b. Flow cytometry confirmations were completed both before and after each time course. (B) A schematic of the BMDM 48-hour time course with staggered time point preparation to optimize the collection window. Four groups (G1-4) of plates each were serum shocked with 12 hour staggering between each so four time points could be processed simultaneously every 2 hours. For 6 total time courses, BMDMs were collected from the bone marrow of a single male PER2::LUC mouse on day 1 and incubated at 37°^C^ for 6 days in Dulbecco’s Modified Eagle Media (DMEM) supplemented with 10% Fetal Bovine Serum (FBS) to full confluence (red). On day 6 media was replaced with Lebovitz supplemented with 10% FBS (yellow) for 24 hours followed by 24 hours in a Leibovitz starve media containing 0% FBS (gray). BMDMs were then shocked for 2 hours in 50% FBS Lebovitz media (green) and assayed for the time course in 10% FBS Leibovitz with luciferin (black) in plates sealed with glass coverslips. (C) A representative luminescence tracing of the tracings that were used to confirm both the robust rhythmicity of PER2 as well as the synchronization between plates within and between time course experiments. Each line corresponds to a plate of macrophages.

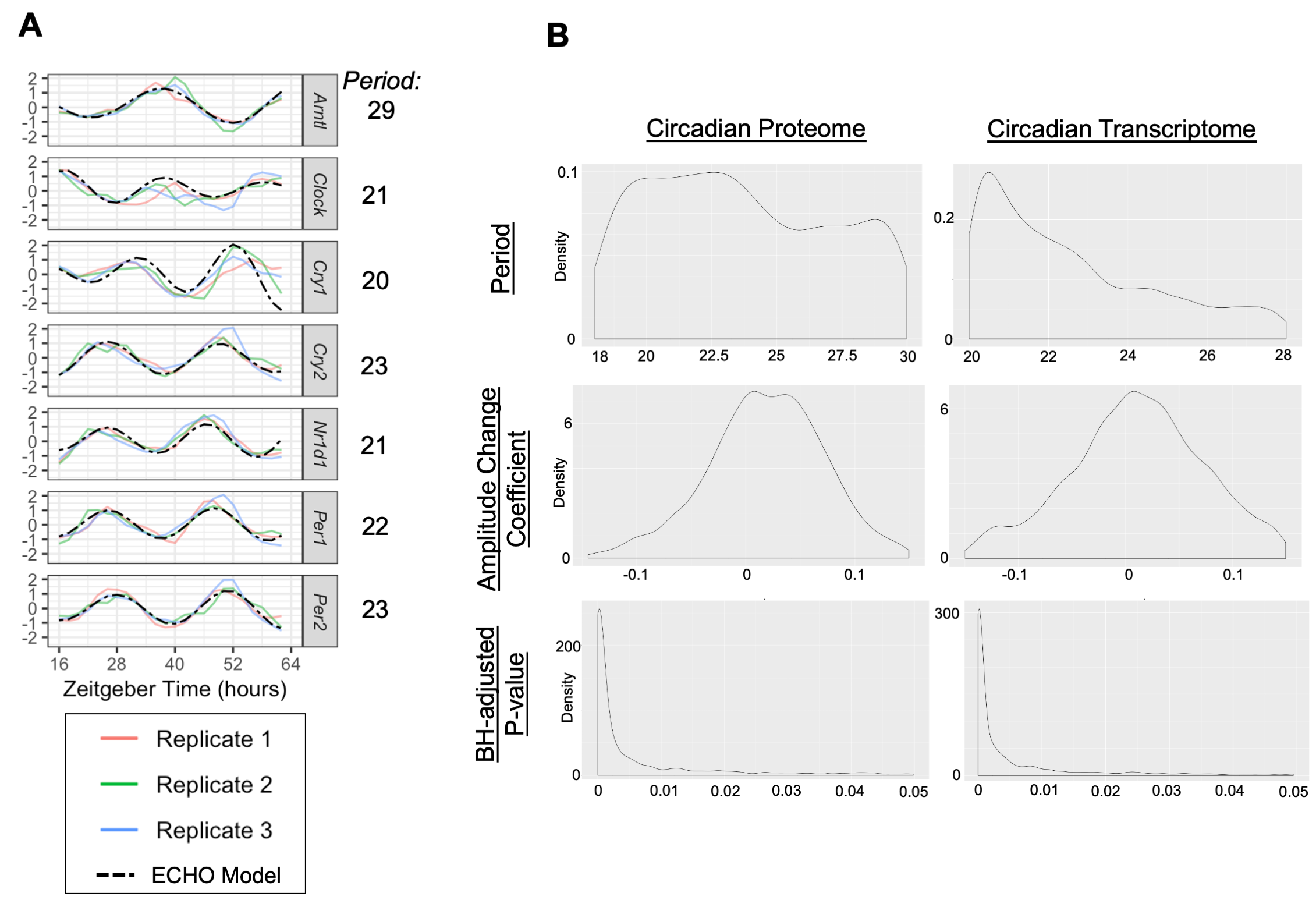

**Supplemental Figure 2. ECHO Modeling of Circadian Oscillations.**

(A) Graphs of the modeled oscillator determined by ECHO for each of the clock gene transcripts plotted with corresponding replicate data. Best fitting model periods were calculated by ECHO ranged from 20-29 hours. (B) Density plots for several ECHO model parameters for the genes identified as circadian in the proteome (left panels) and transcriptome (right panels). X-axis represents parameter corresponding to panel row name.

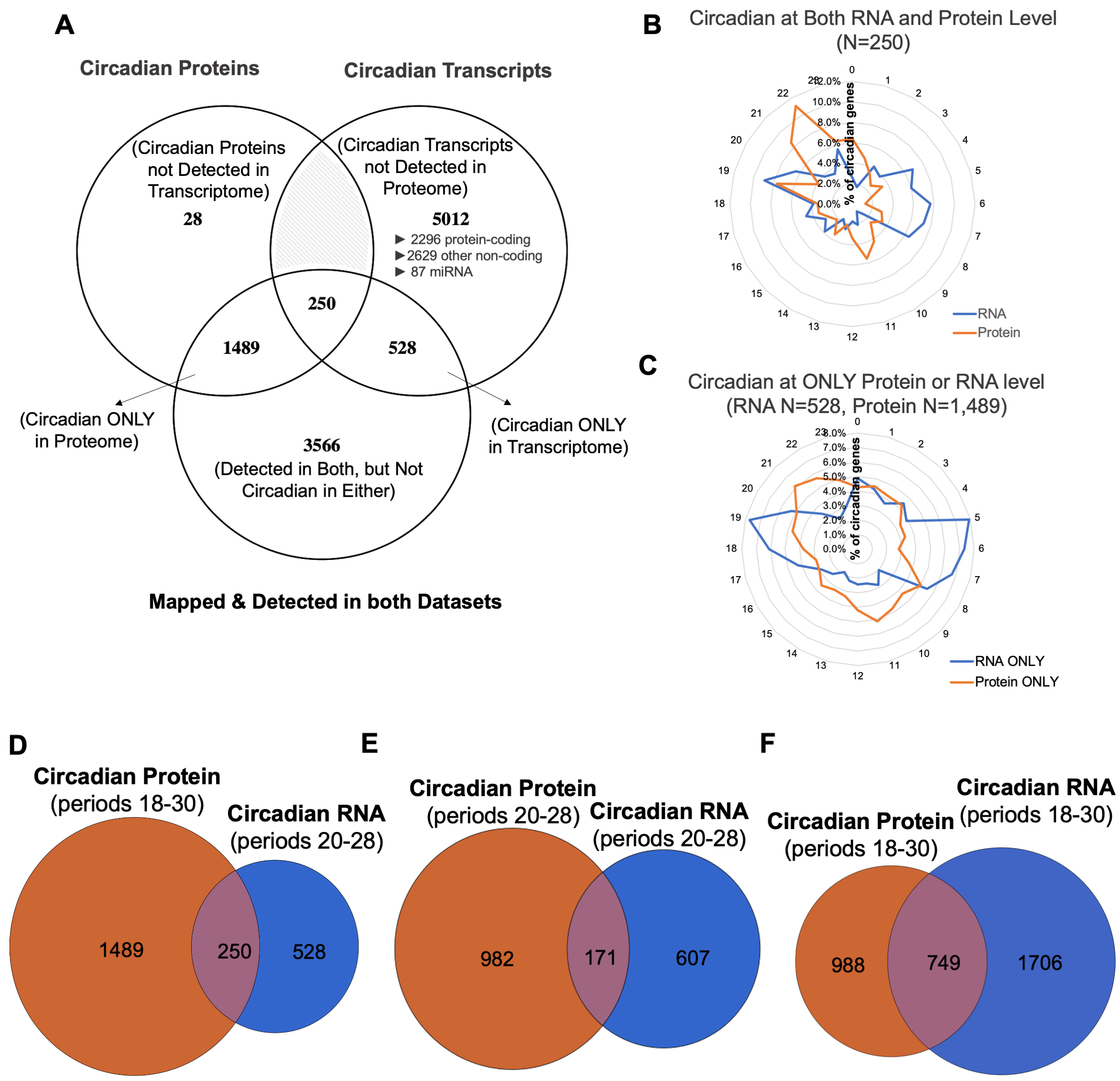

**Supplemental Figure 3. Transcriptome and proteome analysis demonstrate significant discordance.**

(A) Venn diagrams displaying the overlap/discordance between circadian RNAs and circadian proteins. Datasets were subset to compare which proteins and transcripts were detected in both datasets before being compared to each other to determine which proteins were rhythmic in protein, RNA, or both. The resultant oscillating transcripts and proteins were plotted on a clock style graph by Circadian time (CT) (transcripts = blue, proteins = orange) (B and C). Venn diagrams displaying the overlap between circadian RNA (blue) vs Protein (orange) with a protein period cut off of 18-30 and transcript period cutoff of 20-28 (C). More stringent cutoffs of 20-28 for both (E) and less stringent cutoffs of 18-30 for both (F) were also visualized.

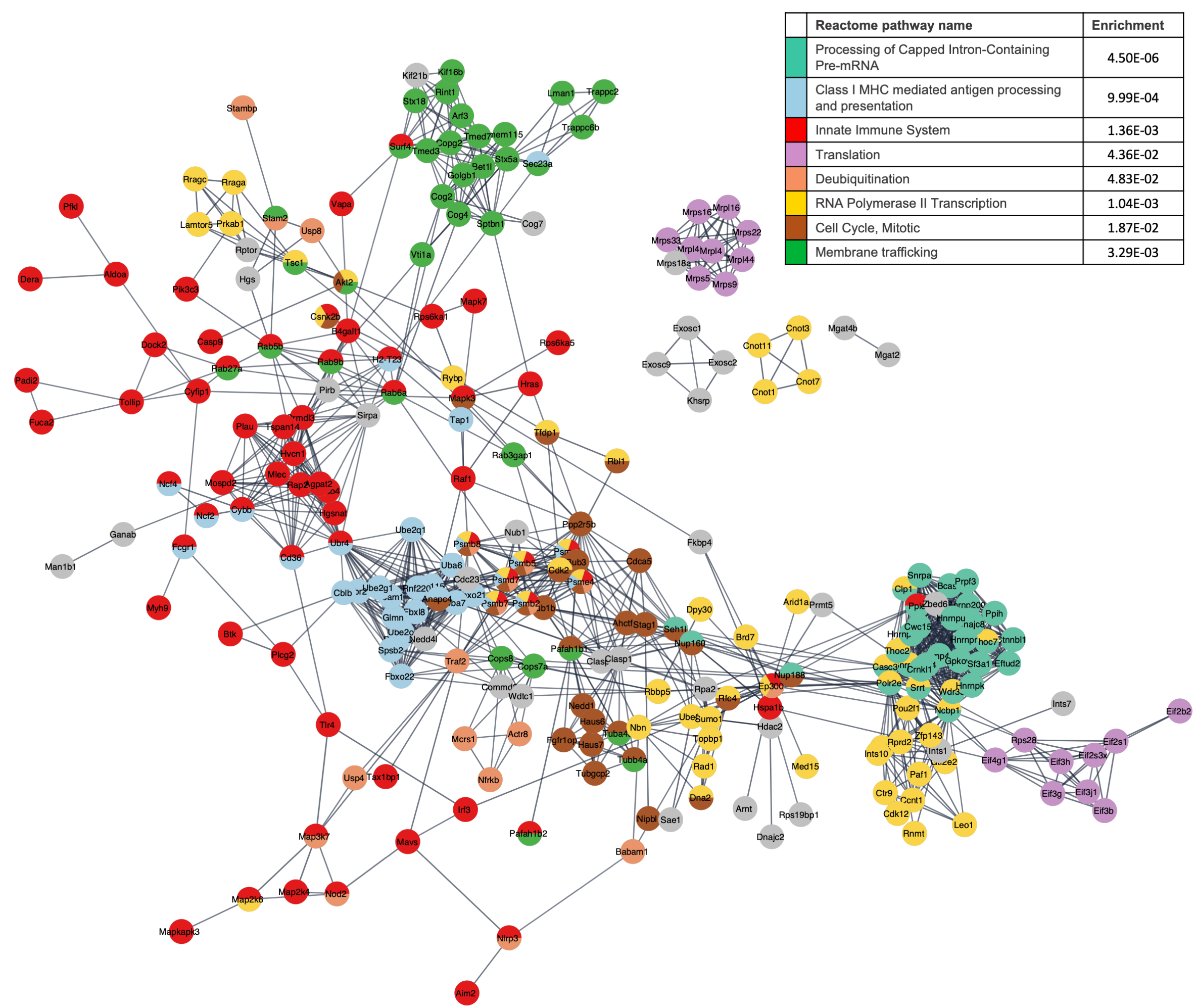

**Supplemental Figure 4. Protein-Protein Interaction Network of the Late Active Wave.**

A high-resolution overview of the late active wave (CT15-3) protein-protein interaction network with individual gene symbols displayed on each node. Strings shown are high confidence (>.9) interactions and node color corresponds to a gene’s enriched reactome category. Grey coloring represents genes from enriched reactome categories that did not represent a large proportion of the protein-protein interactome or belonged to redundant categories.

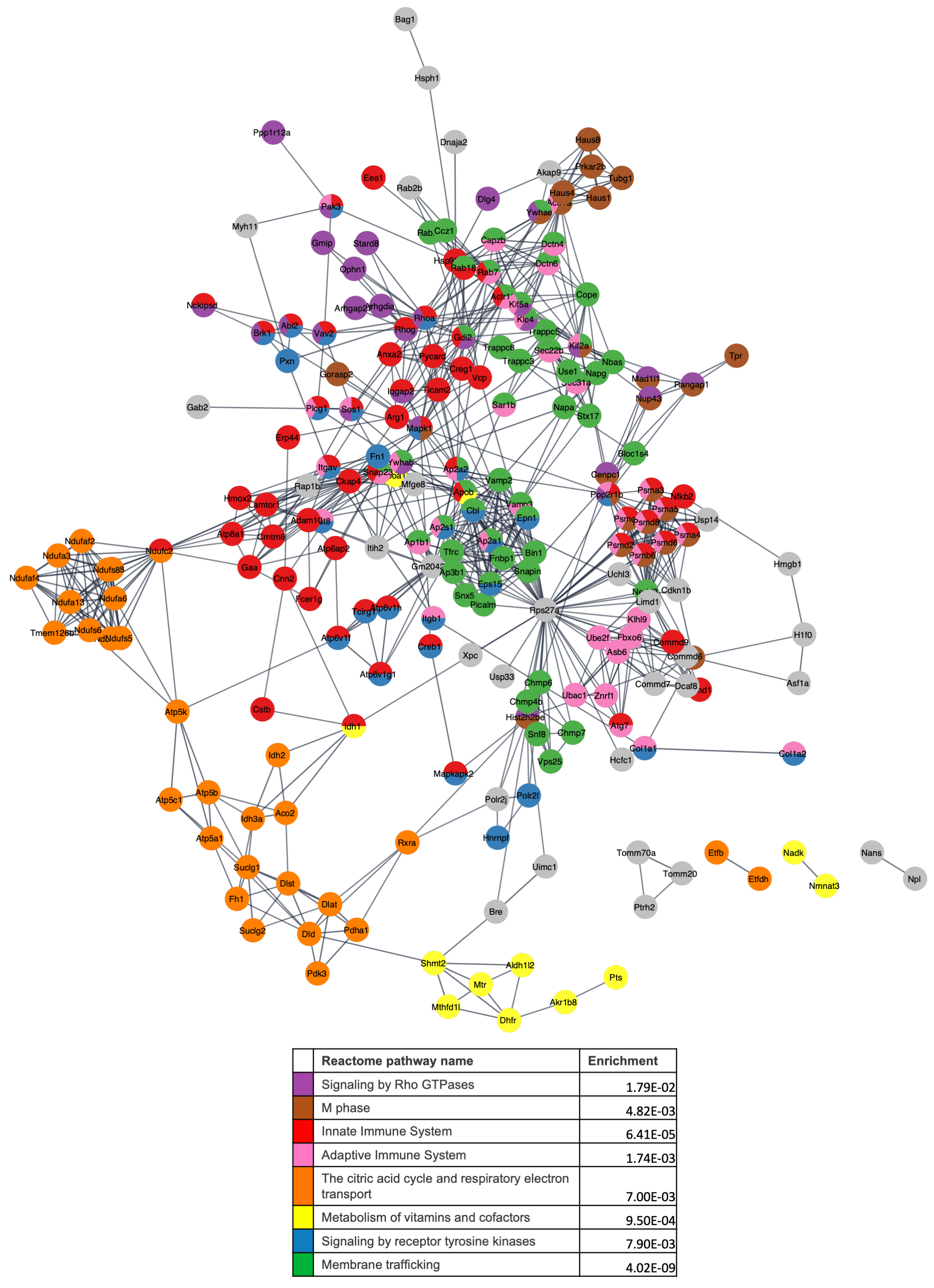

**Supplemental Figure 5. Protein-protein interaction network of the late inactive wave.**

A high-resolution overview of the late inactive wave (CT3-15) protein-protein interaction network with individual gene symbols displayed on each node. Strings shown are high confidence (>.9) interactions and node color corresponds to a gene’s enriched reactome category. Grey coloring represents genes from enriched reactome categories that did not represent a large proportion of the protein-protein interactome or belonged to redundant categories.

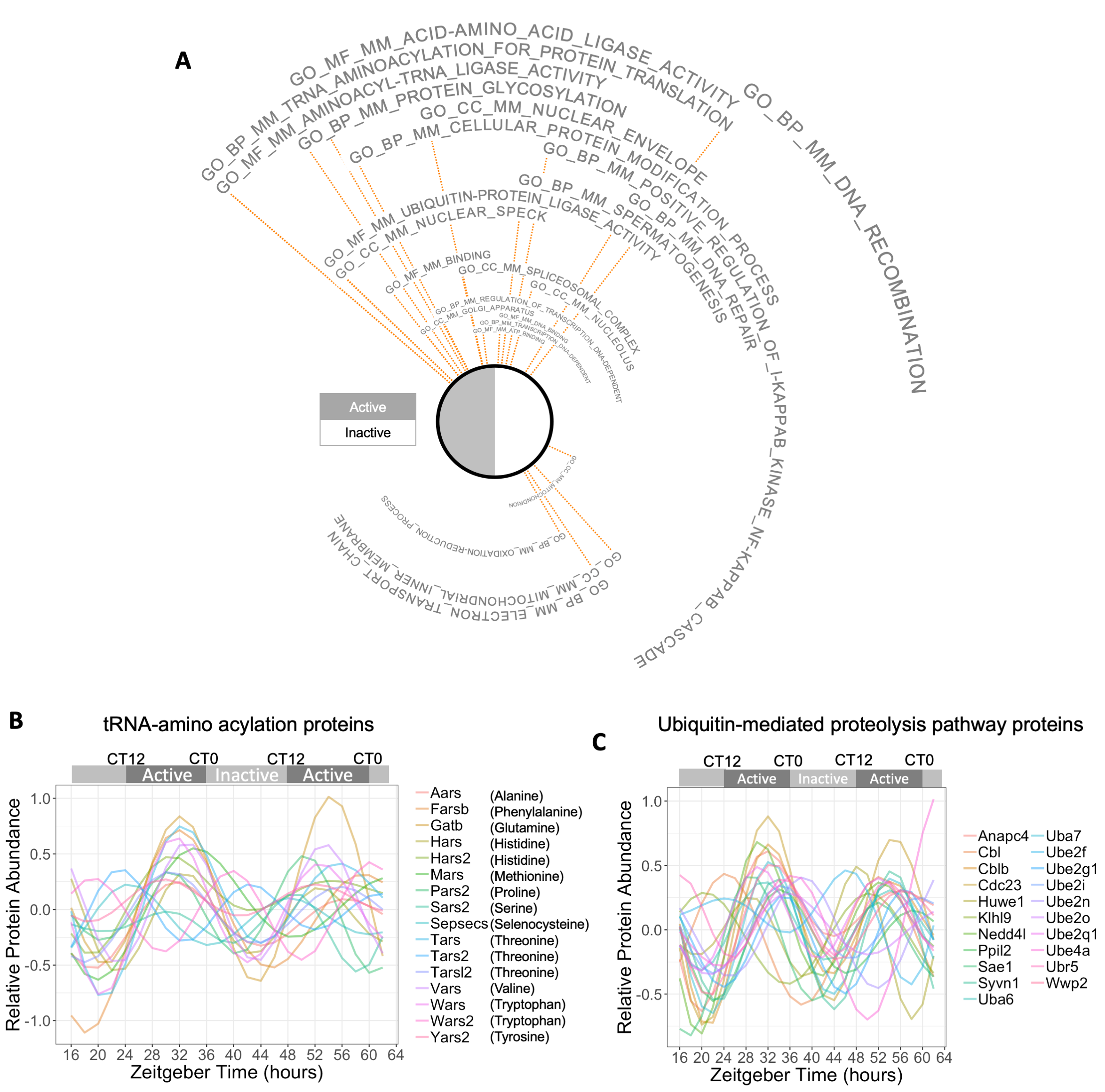
**Supplemental Figure 6. Phase Set Enrichment Analysis Reveals Translational and Degradative Pathways Peak in the Late Active Wave**

(A) Phase Set Enrichment Analysis (PSEA) of the circadian proteome peaking using mouse gene sets curated by Gene Set Knowledgebase (GSKB). The size of text corresponds to the statistical overrepresentation assuming uniform distribution and placement indicates the circadian time of day that the enriched gene set was calculated to occur. (B) Individual modeled fits of all circadian proteins with aminoacyl-ligase activity. The amino acid ligated by each gene is shown in the legend. (C) Individual modeled fits of all circadian proteins in the ubiquitin-proteasome pathway.

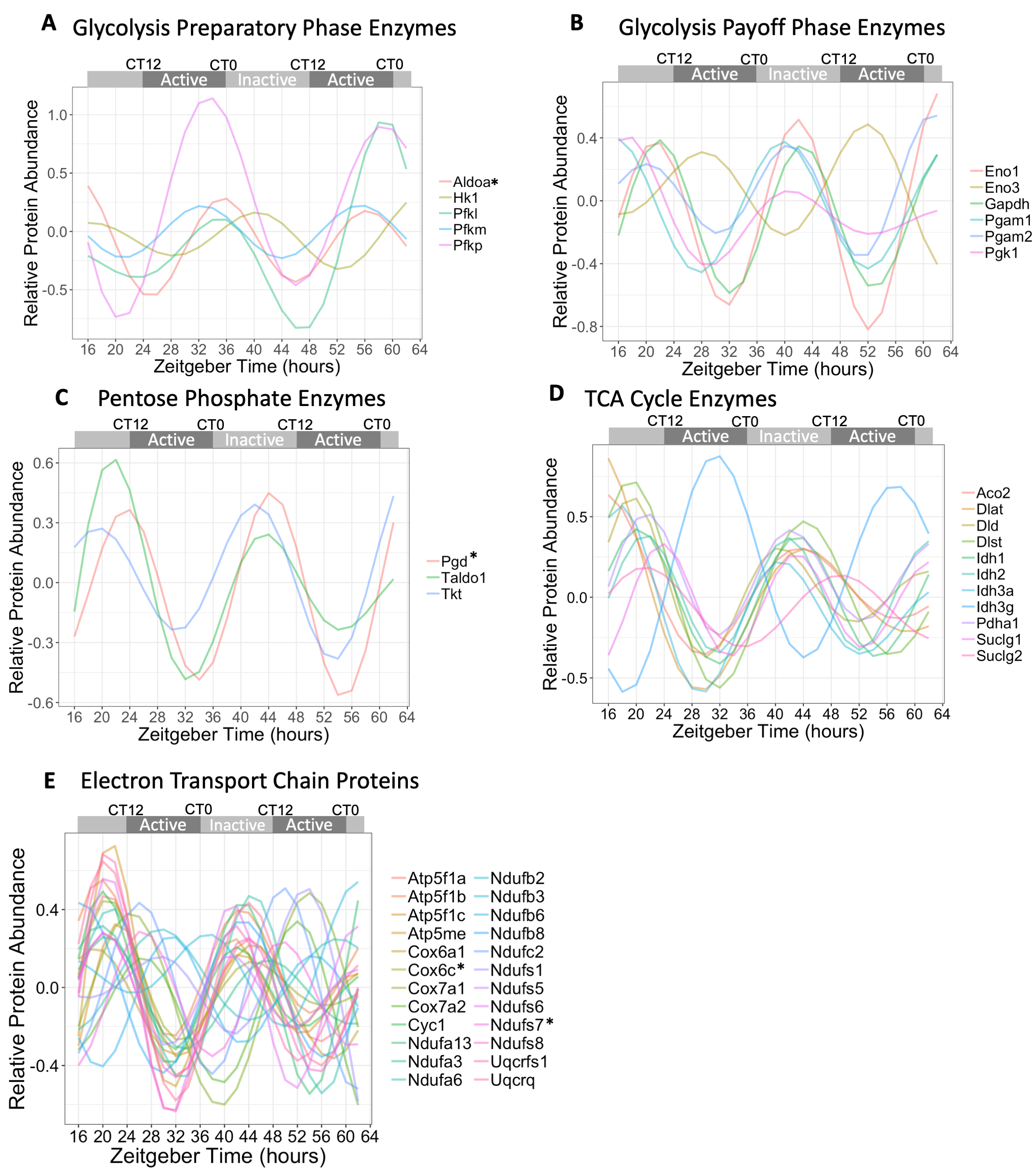
**Supplemental Figure 7. Circadian Proteins in Central Metabolic Pathways**

(A) Individual modeled fits of all circadian proteins in the glycolysis preparatory phase. Asterisk denotes protein also identified as circadian at mRNA level. (B) Individual modeled fits of all circadian proteins in the glycolysis payoff phase. (C) Individual modeled fits of all circadian proteins in the Pentose Phosphate Pathway. Asterisk denotes protein also identified as circadian at mRNA level. (D) Individual modeled fits of all circadian proteins in the TCA cycle. (E) Individual modeled fits of all circadian proteins in the glycolysis payoff phase. (C) Individual modeled fits of all circadian proteins in the Electron Transport Chain. Asterisk denotes protein also identified as circadian at mRNA level.

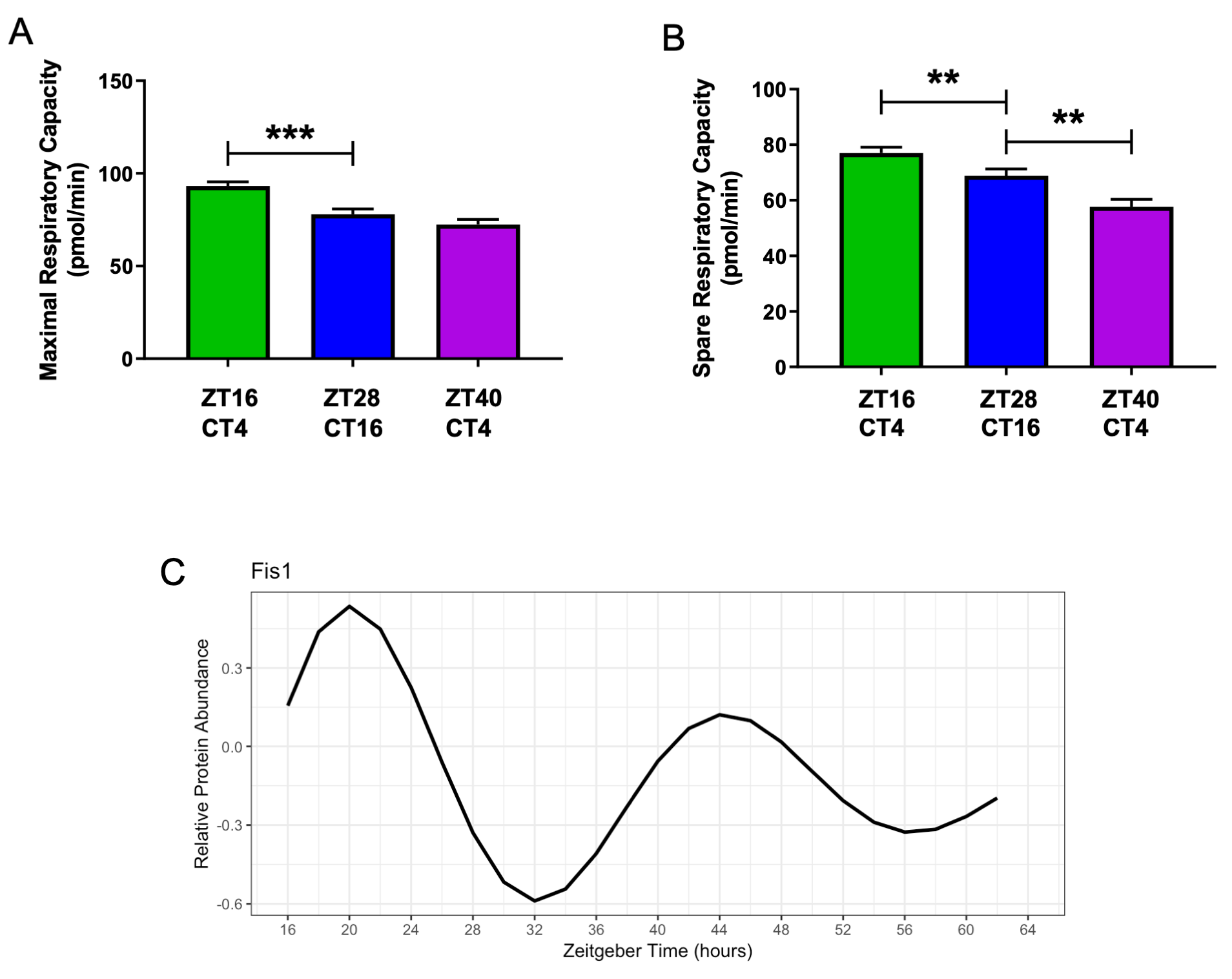
**Supplemental Figure 8. Mitochondrial Maximal Respiration and Spare Respiratory Capacity Decreases over Time.** SEAHORSE-derived measurements for the rate of (A) Maximal respiration and (B) spare respiratory capacity. (C) ECHO modeled fit of Fis1, another identified circadian protein potentially involved in mitochondrial fission.

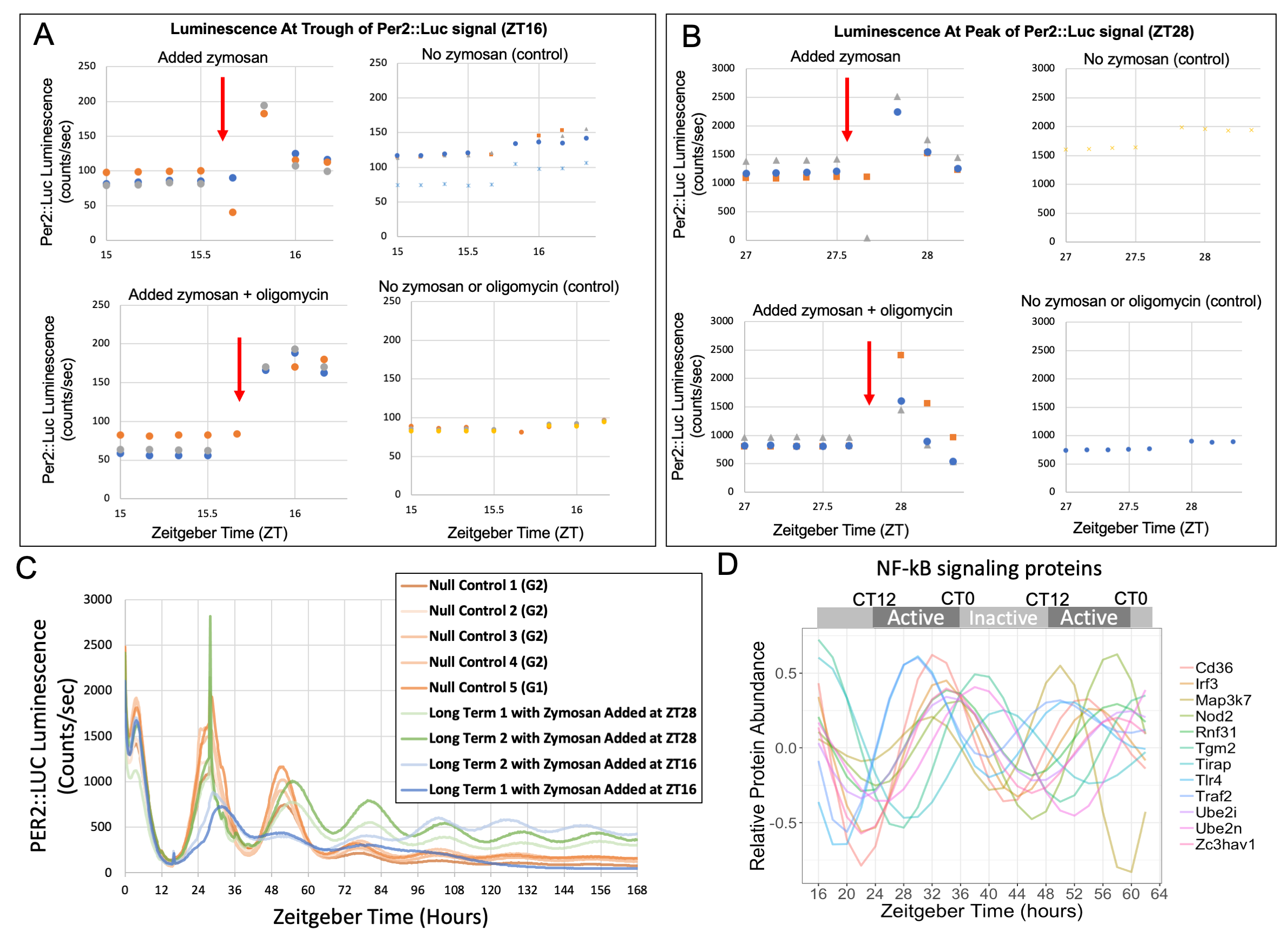

**Supplemental Figure 9. Impact of Zymosan/Oligomycin on the Oscillation of Per2.**

Luminescence counts representing Per2 levels immediately following the addition of Zymosan or Zymosan/Oligomycin at (A) ZT16 and (B) ZT28 demonstrates that there is a slight immediate effect of the core rhythms of the clock, but is proportional to the Per2 level prior to zymosan addition and generally returns to baseline within the time of the 30 minute assay. (C) Long term tracings following zymosan addition shows Per2 oscillations continue despite addition of zymosan, although with phase-shift and amplitude changes depending on time of exposure to Zymosan. (D) Individual modeled fits of all circadian proteins in the NF-kB signaling pathway.

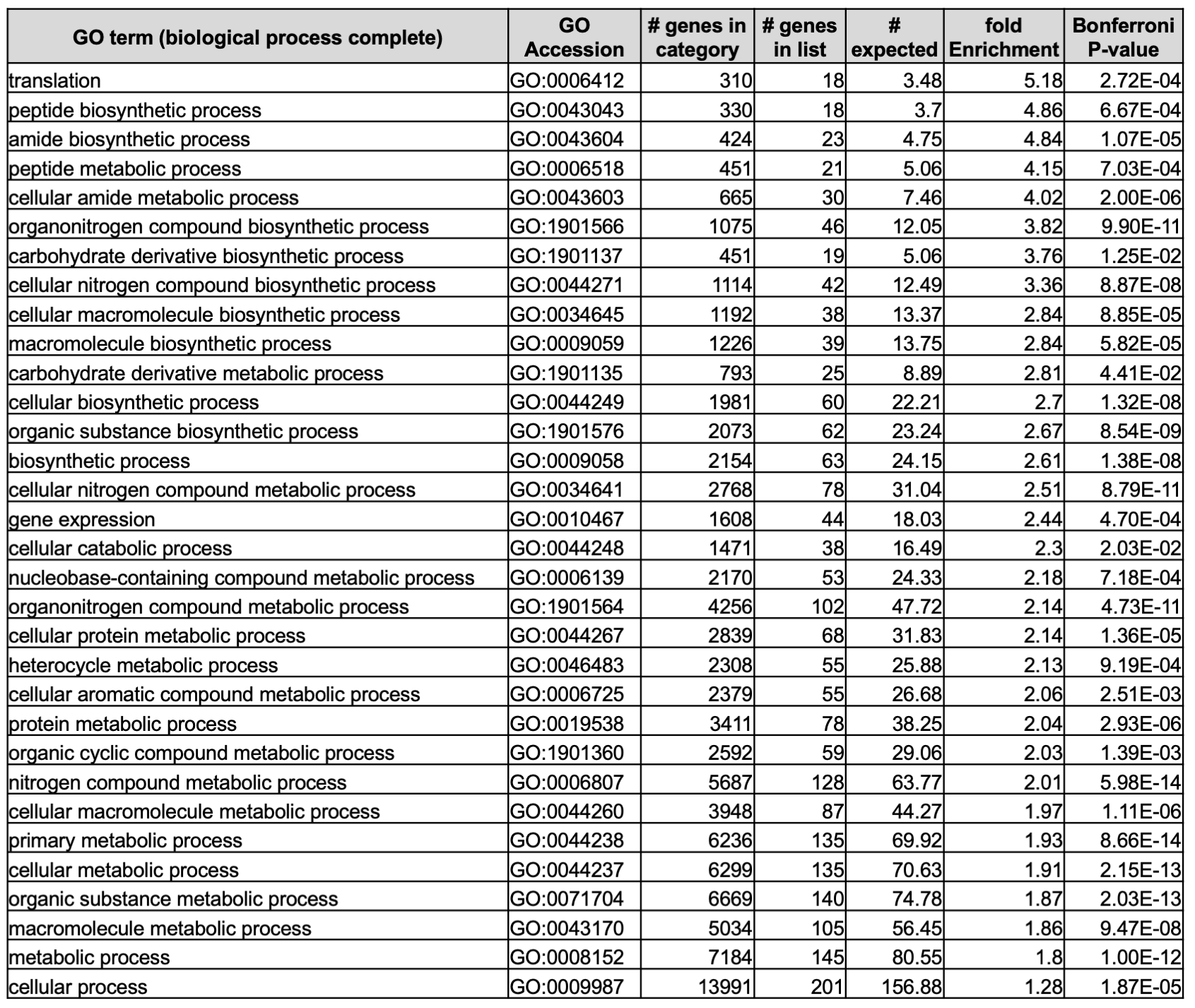
SUPPLEMENTAL TABLES

**Supplemental Table 1: Biological process category enrichments for the 250 genes that are circadian at both the mRNA and protein levels**

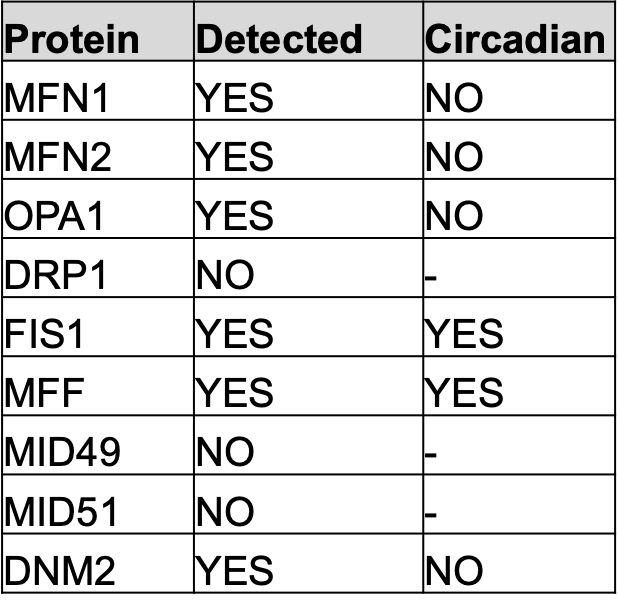

**Supplemental Table 2: Mitochondrial fission/fusion proteins evaluated for detection and circadian rhythmicity in proteome dataset.**

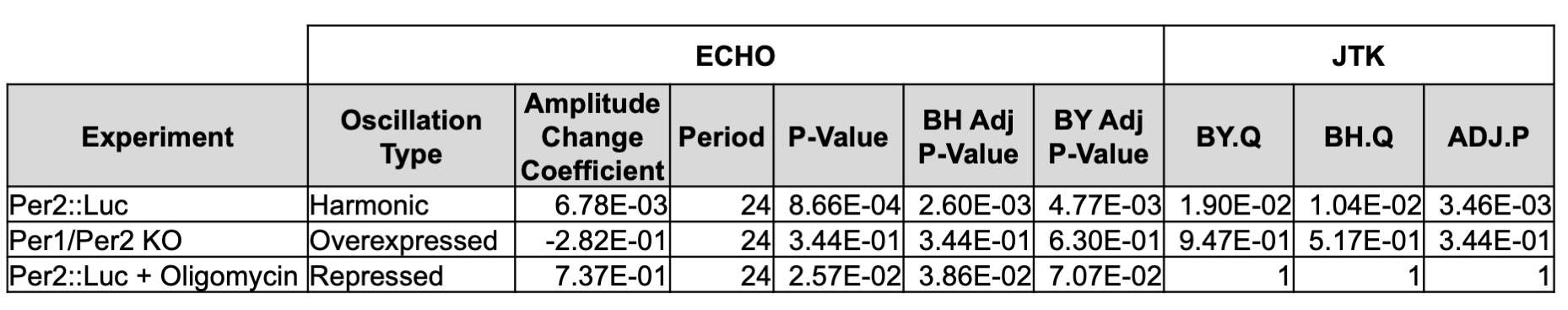
**Supplemental Table 3: ECHO model parameters for phagocytosis assay data.** ECHO was run with smoothing and JTK options.
